# Intrinsic and extrinsic determinants of conditional localization of Mms6 to magnetosome organelles in *Magnetospirillum magneticum* AMB-1

**DOI:** 10.1101/2024.01.08.574735

**Authors:** Carson D. Bickley, Arash Komeili

## Abstract

Magnetotactic bacteria are a diverse group of microbes that use magnetic particles housed within intracellular lipid-bounded magnetosome organelles to guide navigation along geomagnetic fields. Development of magnetosomes and their magnetic crystals in *Magnetospirillum magneticum* AMB-1 requires the coordinated action of numerous proteins. Most proteins are thought to localize to magnetosomes during the initial stages of organelle biogenesis regardless of environmental conditions. However, magnetite-shaping protein Mms6 is only found in magnetosomes that contain magnetic particles, suggesting that it might conditionally localize after the formation of magnetosome membranes. The mechanisms for this unusual mode of localization to magnetosomes are unclear. Here, using pulse-chase labeling, we show that Mms6 translated under non-biomineralization conditions translocates to pre-formed magnetosomes when cells are shifted to biomineralizing conditions. Genes essential for magnetite production, namely *mamE, mamM,* and *mamO* are necessary for Mms6 localization, whereas *mamN* inhibits Mms6 localization. MamD localization is also investigated and found to be controlled by similar cellular factors. The membrane localization of Mms6 is dependent on a glycine-leucine repeat region, while the N-terminal domain of Mms6 is necessary for retention in the cytosol and impacts conditional localization to magnetosomes. The N-terminal domain is also sufficient to impart conditional magnetosome localization to MmsF, altering its native constitutive magnetosome localization. Our work illuminates an alternative mode of protein localization to magnetosomes in which Mms6 and MamD are kept in the cytosol by MamN until biomineralization initiates, whereupon they translocate into magnetosome membranes to control the development of growing magnetite crystals.

**Importance:** Magnetotactic bacteria (MTB) are a diverse group of bacteria that form magnetic nanoparticles surrounded by membranous organelles. MTB are widespread and serve as a model for bacterial organelle formation and biomineralization. Magnetosomes require a specific cohort of proteins to enable magnetite formation, but how those proteins are localized to magnetosome membranes is unclear. Here, we investigate protein localization using pulse-chase microscopy and find a system of protein coordination dependent on biomineralization permissible conditions. In addition, our findings highlight a protein domain that alters the localization behavior of magnetosome proteins. Utilization of this protein domain may provide a synthetic route for conditional functionalization of magnetosomes for biotechnological applications.

## Introduction

The formation of lipid membrane-bounded organelles in eukaryotes is a complex task requiring the activity and coordination of many proteins. Several bacteria also create organelles and must localize specific proteins to developing compartments. One of the best-studied bacterial organelles is the magnetosome, produced by magnetotactic bacteria (MTB) (1). MTB are a diverse set of Gram-negative bacteria that synthesize the magnetic minerals magnetite (Fe_3_O_4_) or greigite (Fe_3_S_4_) (2,3,4). The magnetic crystals are produced within a lipid membrane to form a magnetosome, and magnetosomes are aligned into one or more chains across the cell to create a stable magnetic dipole (5,6). MTB use magnetosome chains to align themselves with the Earth’s magnetic field, allowing for a more efficient search for preferred positions in the oxic-anoxic transition zone (1,2).

In the model organisms *Magnetospirillum magneticum* AMB-1 and *Magnetospirillum gryphiswaldense* MSR-1, magnetosome biogenesis is performed mainly by proteins encoded by the magnetosome gene island (MAI) (7). Magnetosome genes identified in the MAI are organized into four clusters (*mamAB, mamGFDC, mamXY,* and *mms6*) which are necessary and sufficient for magnetosome formation (8). Many MAI proteins localize specifically to magnetosome membranes and are depleted in other cellular membranes (9,10,11,12,13). Little is known about how proteins are sorted to magnetosomes – magnetosome proteins lack a universal signal peptide (14,15) – but it is thought that they may aggregate on the inner membrane at magnetosome development sites in the early stages of magnetosome membrane invagination (15,16). Aggregated proteins would therefore be concentrated into magnetosome membranes as compartments form.

In contrast, recent evidence suggests that the biomineralization protein Mms6 may localize to magnetosomes after membrane formation is complete (16). Mms6 was originally isolated in a proteomic search for proteins tightly bound to magnetite crystals of AMB-1. It is predicted to have a transmembrane region and has been identified in enriched magnetosome membranes (13). The Mms6 N-terminus is thought to either associate with the magnetosome membrane surface or translocate through the membrane, while the C-terminal region contacts the magnetite (17). Accordingly, *in vitro* magnetite synthesis in the presence of the C-terminal 6 kDa region of Mms6 (18,19), or even the acidic peptide contained within it (19,20), results in cubo-octahedral crystals resembling those produced *in* vivo. Mutations in *mms6* result in the formation of smaller, misshapen crystals, further indicating a role in magnetite crystal shaping (19,21,22). Magnetite-binding activity has been suggested to be necessary for Mms6 localization (23). A study by Arakaki et al. in AMB-1 used correlated transmission electron microscopy and fluorescence microscopy to show that Mms6 only localizes to magnetosome membranes that contain magnetite (16). When AMB-1 is grown under oxic conditions that do not permit biomineralization, Mms6-GFP localizes diffusely throughout the cell, although whether Mms6-GFP localized either in the inner membrane or cytoplasm was unclear (16). When conditions are changed to permit biomineralization, Mms6 is localized to magnetosome membranes in as few as 2 h (16). In contrast, many proteins such as fellow crystal maturation proteins MamC (also known as Mms13) and MmsF were found to localize to magnetosome membranes even under oxic conditions that prevent crystal formation (16).

While previous work has determined an unusual localization mode for Mms6, the dynamics of the process, as well as the extrinsic and intrinsic molecular factors governing it, have remained obscure. Therefore, we combined pulse-chase analyses, imaging, and genetic analyses to define the process of Mms6 localization at a molecular level. We show that upon a shift into biomineralization-permissible conditions (BPC), pre-translated Mms6 relocalizes from the cytosol to pre-formed magnetosome membranes, displaying a surprising localization behavior for a protein containing a transmembrane domain. We also identify three genes, *mamE, mamO,* and *mamM*, that are necessary for Mms6 localization. In contrast, *mamN* is implicated in inhibiting Mms6 localization to empty magnetosomes. We identify MamD as a protein that, like Mms6, conditionally localizes to magnetosomes in the presence of *mamN*, suggesting the mechanisms regulating Mms6 localization also regulate other magnetosome proteins. Additionally, we show that the N-terminal domain of Mms6 is necessary for retention in the cytosol and can impart conditional localization on a heterologous magnetosome protein. We speculate that AMB-1 responds to BPC by sorting pre-folded Mms6 to magnetosome membranes to help shape the developing crystal, exhibiting a more dynamic and complex strategy of regulating biomineralization than previously hypothesized. Exploiting this strategy in synthetic applications could allow fine-tuning of biocompatible magnetic nanoparticles, which have wide ranging applications including targeted drug delivery (24), MRI contrast enhancement (25), and magnetic hyperthermia therapy (26).

## Results

### Translated Mms6 relocalizes to pre-existing magnetosomes after the initiation of biomineralization

Mms6-GFP was shown to localize to magnetosome membranes only when conditions favor biomineralization (16). Understanding the dynamics of this process could provide insights into the mechanisms and functional relevance of conditional protein sorting to magnetosome organelles. Previously, Mms6-GFP was imaged after biomineralization was induced in one of two ways, either by moving cells grown under microaerobic conditions to anaerobic conditions, or by adding iron to iron-starved cells (16). Images taken 2 h after iron addition or 8 h following growth in anaerobic conditions revealed that Mms6-GFP had localized to magnetosomes (16). To replicate these results, we first grew cells in iron starvation conditions. Iron was added to cultures to induce biomineralization, and the localization patterns of Mms6-GFP in live cells was grouped into three categories: “Foci”, indicating cells with one or more unaligned fluorescent foci, “Diffuse”, indicating protein diffuse in the cytoplasm, and “Chain”, indicating linear fluorescent patterns consistent with proteins localized to the magnetosome chain. Mms6-GFP was primarily diffuse in the cytosol in cells grown without added iron (Fig. 1A,B). In line with previous results, Mms6-GFP was localized to magnetosome chains in most cells 1-2 h post induction (Fig. 1A,B).

**Figure 1.**
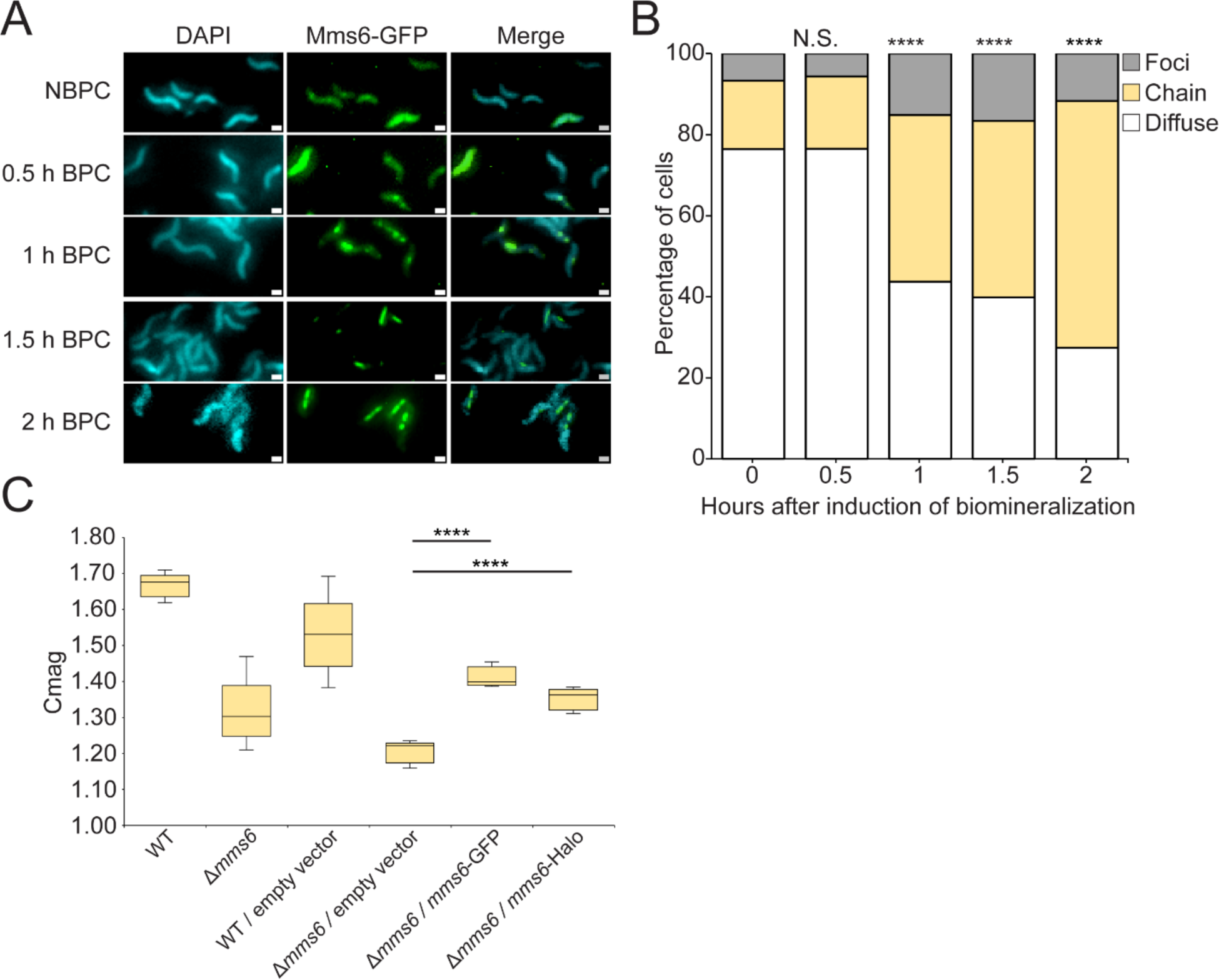
Mms6-GFP localizes to magnetosomes in response to biomineralization permissible conditions. (A) Representative fluorescence microscopy images of *Δmms6* expressing Mms6-GFP shown in green. DAPI is shown in blue. Scale bars = 1 µm. (B) Blind quantification of localization patterns of Mms6-GFP during biomineralization time course *in vivo*. Cells were categorized by localization pattern, and y-axis represents percentage of cells with Mms6-GFP displaying given localization pattern out of total labeled cells. *P* values were calculated by Chi-squared test of independence comparing given dataset to time 0 (N.S. no significant difference *P* > .01) (**** *P* < 10^-5^). NBPC *n* = 686 cells, 0.5 h *n* = 1147 cells, 1 h *n* = 507 cells, 1.5 h *n* = 1013 cells, 2 h *n* = 1255 cells. (C) Coefficient of magnetism (Cmag) of strains. *P* values were calculated by Mann-Whitney U test comparing given dataset to Δ*mms6* / empty vector (**** *P* < 10^-5^).

The change in Mms6-GFP localization after iron addition could be the result of two different phenomena. In one model, newly synthesized Mms6 in biomineralization-permissible conditions (BPC) localizes to magnetosomes, while pre-existing Mms6 is diluted by growth and protein turnover. Alternatively, pre-existing Mms6 synthesized under non-biomineralization permissible conditions (NBPC) may relocalize to magnetosomes upon a change in conditions. To differentiate between these possibilities, we used the HaloTag protein fusion tag, which covalently and irreversibly binds to fluorescent ligands, allowing the tracking of a specific protein pool (27). Mms6-Halo expressed in a *Δmms6* background partially restored the cellular magnetic response, assayed by determining the Coefficient of magnetism (Cmag), a measurement dependent on the differential scattering of light by cells moved into different orientations by an external magnetic field (Fig. 1C).

Using HaloTag, we tracked a pool of Mms6-Halo synthesized before iron was added to non-biomineralizing cells. To determine if the old pool of Mms6-Halo relocalized to magnetosomes or if new protein synthesis was necessary, we performed a pulse-chase experiment. Briefly, AMB-1 cultures grown and passaged under iron starvation conditions were labeled with fluorescent ligand JF549. Then, biomineralization was induced and cells were grown for several hours. To ensure that a representative sample of cells was tracked during the pulse-chase experiment, the percentage of cells containing fluorescent protein was calculated throughout the experiment. The percentage of cells containing fluorescent protein decreased only slightly, from 73% to 68% over 2 h of incubation, suggesting that similar representative samples were captured at each timepoint (Fig. 2A). One confounding factor with the pulse chase experiment could be incomplete saturation of HaloTag with fluorescent ligand. To measure label saturation of HaloTag, cells from 10 ml cultures were incubated with a pulse ligand for 1 h, washed, supplemented with bacteriostatic antibiotics (700 µg/ml kanamycin and 400 µg/ml chloramphenicol) to prevent new protein synthesis, incubated for 30 min, and finally incubated with a chase ligand containing a different fluorophore for 1 h. Without antibiotic treatment, similar percentages of cells were labelled with the pulse (82%) and chase (73%) ligands. When treated with antibiotics, 84% of cells were labeled with the pulse ligand, whereas only 27% were labeled with chase ligand, indicating that the pulse ligand conditions saturates most Mms6-Halo proteins (Fig. 2B). Culture growth and magnetic response were tracked during the time course. Cell growth measured by OD_400_ increased steadily after iron addition (black arrow) from an average of 0.050 to 0.188 9 h later, showing that cells remained healthy during the experiment (Fig. 2C). Cmag increased steadily after addition of iron from a starting non-magnetic Cmag of 1.0 and reaching a magnetic Cmag of 1.5 6 h after iron addition (Fig. 2D). Before induction of biomineralization, Mms6-Halo localized diffusely in the cytoplasm in the majority of cells (Fig. 2E,F). In contrast, 2 h after iron addition, the old pool of Mms6-Halo had relocalized to the magnetosome chain in most cells (Fig. 2E,F). These results indicate that Mms6-Halo synthesized under NBPC can relocalize to magnetosomes after the induction of biomineralization.

**Figure 2.**
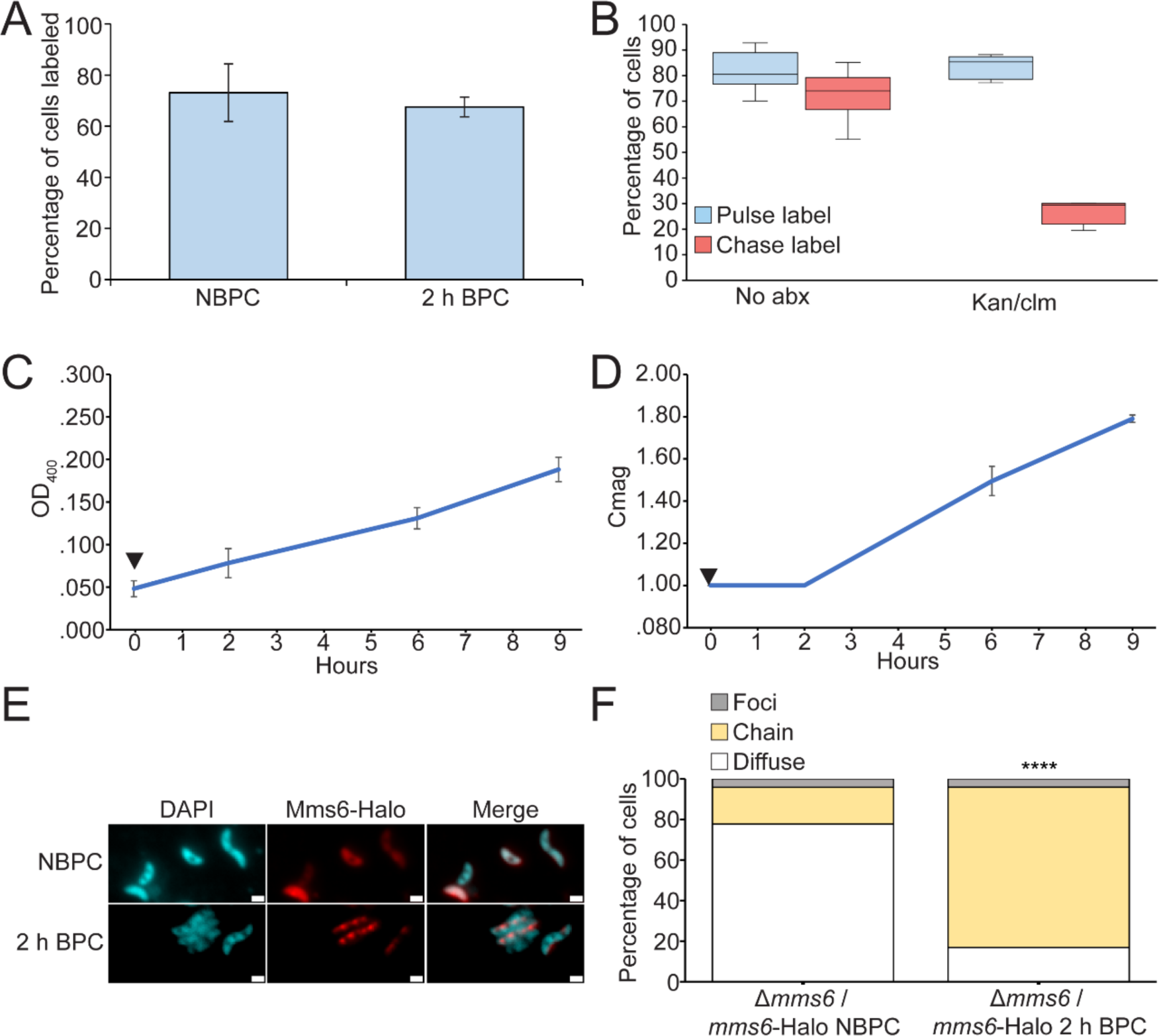
Pre-translated Mms6-Halo is relocalized from the cytoplasm to magnetosomes in response to biomineralization permissible conditions. (A) Percentage of cells labeled with Mms6-Halo fluorescence before and during relocalization time course. NBPC *n* = 12528 DAPI labeled cells, 2 h BPC *n* = 4268 DAPI labeled cells. (B) *Δmms6* / *mms6*-Halo cells were incubated with HaloTag pulse and chase ligands to test HaloTag saturation with and without 700 µg/ml kanamycin and 400 µg/ml chloramphenicol to prevent the synthesis of new Mms6-Halo. (C) OD_400_ of 9 cultures of *Δmms6* / *mms6-Halo* grown initially under iron starvation conditions and then given iron to allow biomineralization (black arrow). (D) Coefficient of magnetic response of cultures over time course. (E) Mms6-Halo with J549 ligand shown in red and DAPI shown in blue. (F) Blind quantification of localization patterns of Mms6-Halo during biomineralization time course *in vivo*. Cells were categorized by localization pattern as above. *P* values were calculated by chi-squared test of independence comparing given dataset to NBPC sample (**** *P* < 10^-5^). NBPC *n* = 8432 labeled cells, BPC *n* = 2810 cells.

To confirm that Mms6-Halo relocalization does not require new protein synthesis, the pulse chase experiment was repeated with bacteriostatic antibiotics to prevent new protein synthesis (Fig. 3A,B). 700 µg/ml kanamycin and 400 µg/ml chloramphenicol were added simultaneously (white arrow) to half of the cultures 1 h before iron addition (black arrow). The antibiotics slowed cell growth and stopped biomineralization, suggesting they prevented new protein synthesis (Fig. 3A,B). The percentage of cells containing fluorescent protein increased slightly, from 74% to 80%, over 2 h of incubation without antibiotics, and from 74% to 86% with antibiotics, suggesting that similar representative samples were captured at each timepoint (Supplementary Fig. 1). Despite the effects of the antibiotics, Mms6-Halo still relocalized to the magnetosome chain like in cells that did not receive antibiotics (Fig. 3C,D), confirming that no new protein synthesis is needed for Mms6 localization. Example images of cells captured using super resolution structured illumination microscopy are provided in Supplementary Figure 2. The timing of relocalization and the patterns observed are most consistent with relocalization to pre-existing chains rather than relocalization exclusively via new magnetosome synthesis (see Discussion). Thus, cytoplasmic Mms6 can relocalize to pre-existing magnetosomes when biomineralization conditions change, revealing a surprising mode of magnetosome protein localization in AMB-1.

**Figure 3.**
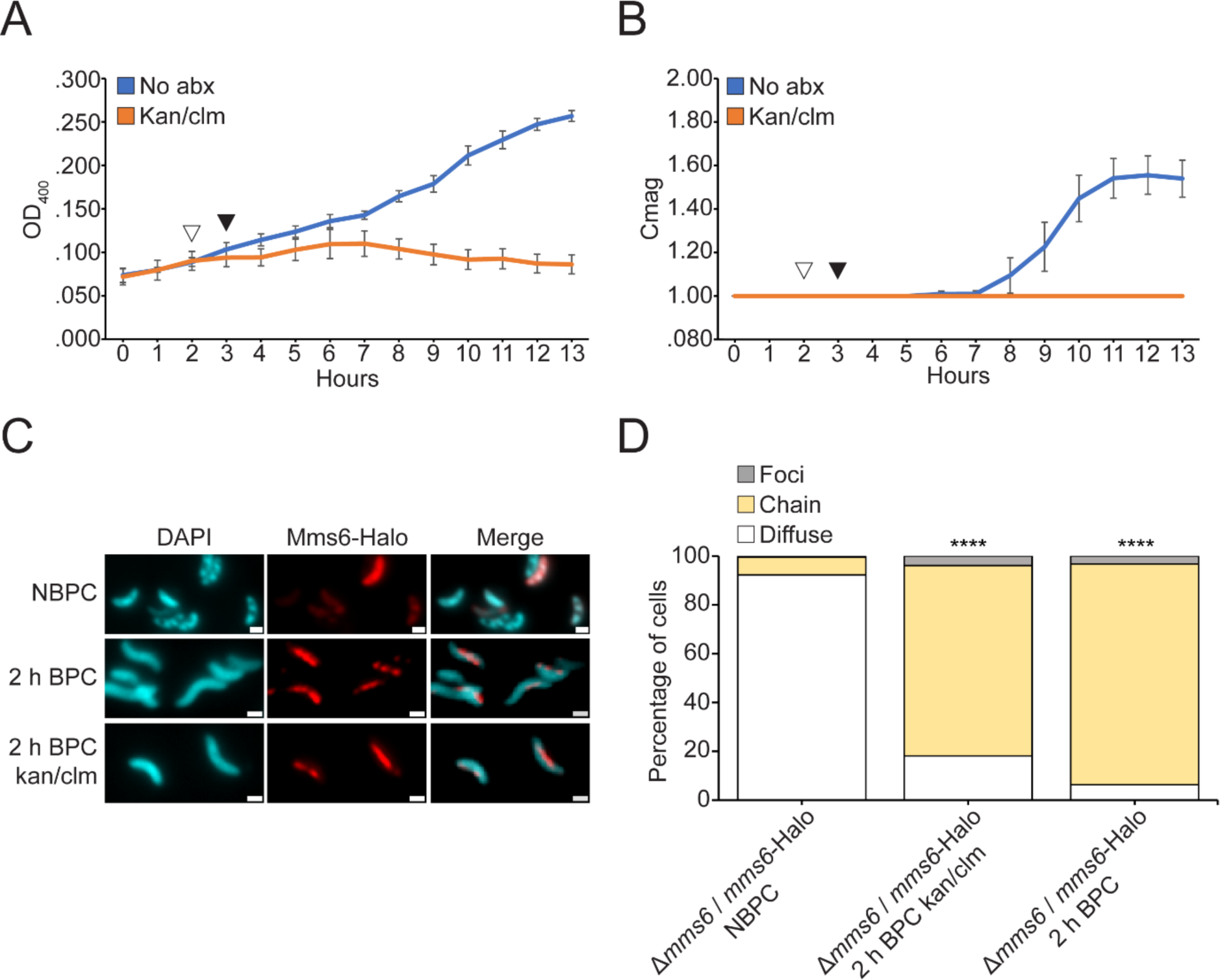
Pre-translated Mms6 relocalizes after iron addition in the absence of new protein synthesis. (A) OD_400_ of *Δmms6* expressing Mms6-Halo grown initially under iron starvation conditions and then given iron to allow biomineralization. 700 µg/ml kanamycin and 400 µg/ml chloramphenicol were added (white arrow) to kan/clm sample to prevent the synthesis of new Mms6 1 h before adding iron (black arrow) to all samples. (B) Coefficient of magnetic response of cultures over time course. (C) Mms6-Halo with J549 ligand shown in red and DAPI shown in blue. (D) Blind quantification of localization patterns of Mms6-GFP during biomineralization time course *in vivo*. Cells were categorized by localization pattern as above. *P* values were calculated by Fisher’s exact test comparing given dataset to NBPC sample (**** *P* < 10^-5^). NBPC *n* = 2112 cells, 2 h BPC kan/clm *n* = 1102 cells, 2 h BPC *n* = 2422 cells.

### MAI proteins are necessary for Mms6 magnetosome localization

Mms6 is a magnetosome-associated protein with a predicted transmembrane domain. Yet, its localization in the absence of magnetite formation is strikingly different from other magnetosome-associated proteins (16). Therefore, we tested whether Mms6 localization is an inherent feature of the protein or requires other magnetosome proteins. In the absence of genes essential to magnetosome formation, many magnetosome-associated proteins are dispersed throughout the cytoplasmic membrane (28). The localization patterns of Mms6-GFP and magnetosome membrane protein GFP-MmsF were examined in mutant cells lacking the MAI and a region outside the MAI that also affects magnetosome positioning, called the magnetotaxis gene islet (MIS) (29). The MAI and MIS contain the majority of known magnetosome proteins in AMB-1, and cells that lack the MAI and MIS are unable to form magnetosome membranes (30). As expected, GFP-MmsF highlights the cell periphery in this mutant, consistent with localization to the inner cell membrane. In contrast, Mms6-GFP (Fig. 4A,B) has a cytosolic localization in ΔMAI ΔMIS cells, even in BPC. Similarly, Mms6-Halo localizes to the cytosol in ΔMAI cells, which are also unable to form magnetosome membranes (Fig. 5A,B). Therefore, Mms6 association with membranes requires either other magnetosome proteins or a previously formed magnetosome membrane.

**Figure 4.**
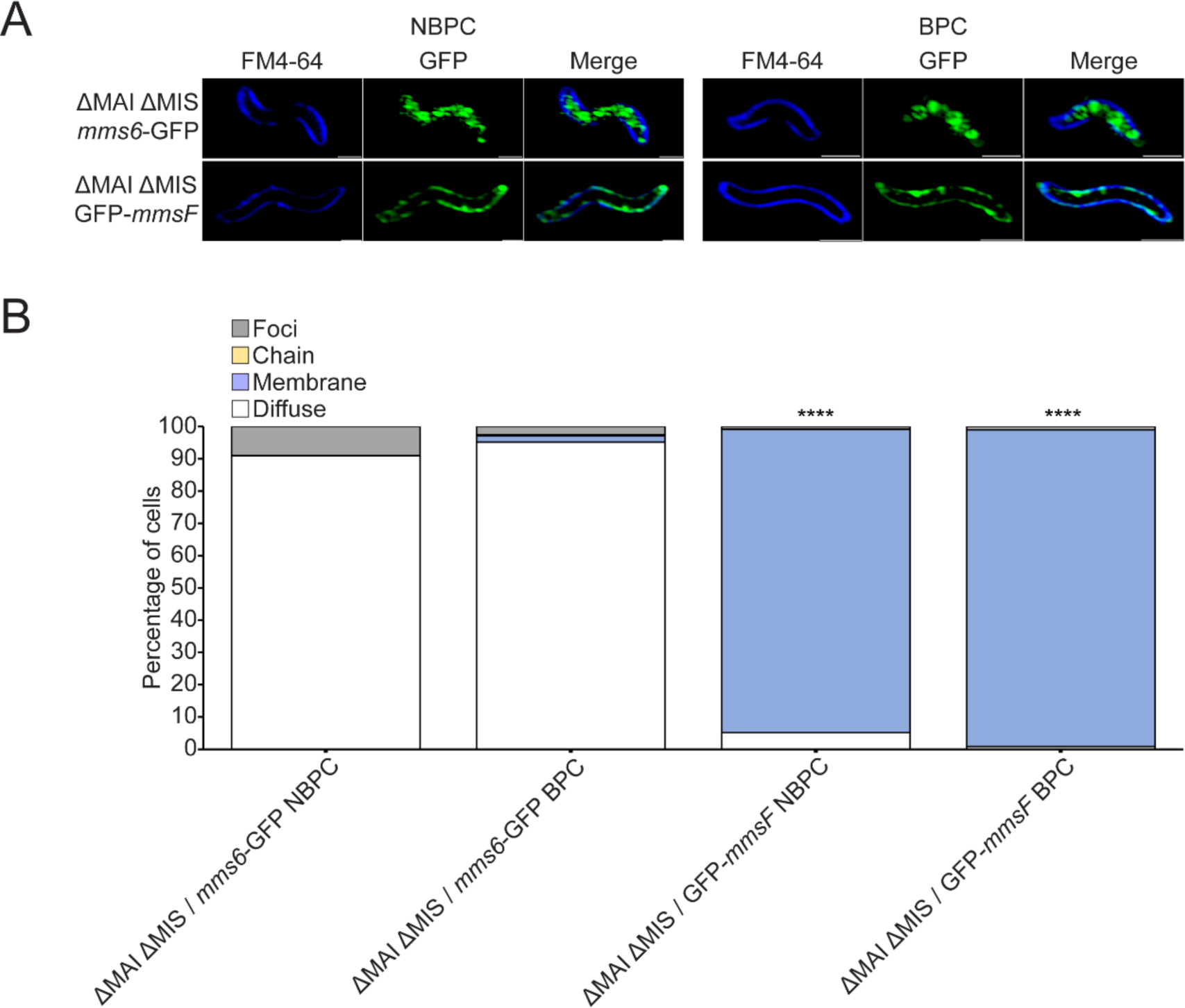
Mms6-GFP is cytoplasmic in the absence of magnetosomes. (A) Representative super resolution 3D Structured Illumination Microscopy (SIM) images of ΔMAI ΔMIS expressing either *mms6*-GFP or GFP-*mmsF* shown in green and membrane stain FM4-64 shown in dark blue. Scale bars = 1 µm. (B) Blind quantification of localization patterns of either *mms6*-GFP or GFP-*mmsF* based on fluorescence microscopy images. Cells were categorized by localization pattern. *P* values were calculated by Fisher’s exact test comparing GFP-*mmsF* dataset to *mms6*-GFP dataset of respective biomineralization condition (N.S. no significant difference *P* > .01) (**** *P* < 10^-5^). ΔMAI ΔMIS / *mms6*-GFP NBPC *n* = 624 cells, ΔMAI ΔMIS / *mms6*-GFP BPC n = 166 cells, ΔMAI ΔMIS / GFP-*mmsF* NBPC n = 588 cells, ΔMAI ΔMIS / GFP-*mmsF* BPC n = 927 cells.

**Figure 5.**
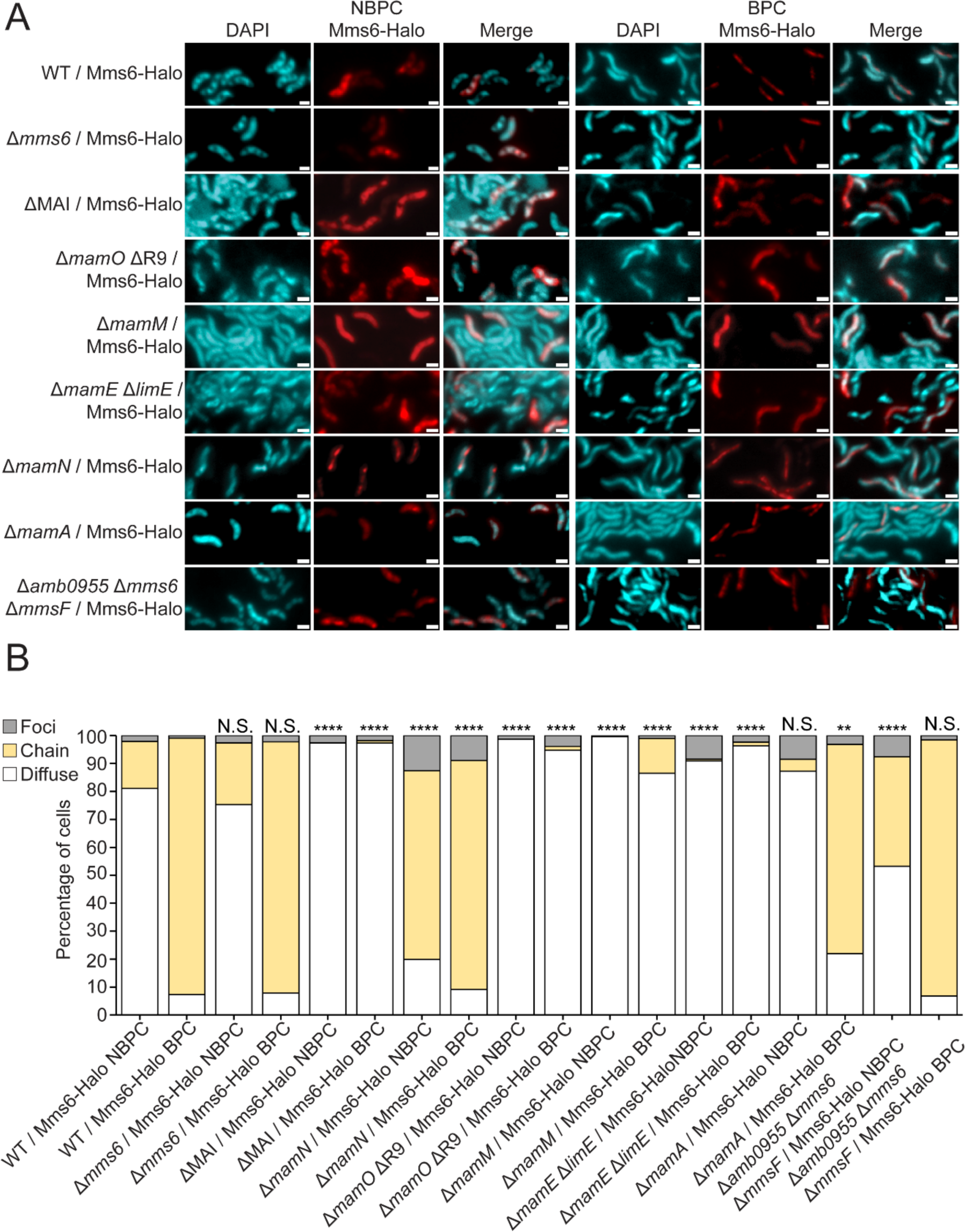
MAI proteins affect Mms6 localization (A) Representative fluorescence microscopy images of AMB-1 with different genetic backgrounds expressing *mms6*-Halo and grown in standard conditions. JF549 HaloTag ligand fluorescence is shown in red and DAPI in blue. Scale bars = 1 µm. (B) Blind quantification of localization patterns of Mms6-Halo. *P* values were calculated by Fisher’s exact test comparing given dataset to WT / *mms6*-Halo (N.S. no significant difference *P* > .01) (**** = *P* < 10^-5^). WT / *mms6*-Halo NBPC *n* = 285 cells, WT / *mms6*-Halo BPC *n* = 436 cells, ΔMAI / *mms6*-Halo NBPC *n* = 496 cells, ΔMAI / *mms6*-Halo BPC *n* = 718 cells, Δ*mamN* / *mms6*-Halo NBPC *n* = 956 cells, Δ*mamN* / *mms6*-Halo BPC *n* = 514 cells, Δ*mamO* ΔR9 / *mms6*-Halo NBPC *n* = 312 cells, Δ*mamO* ΔR9 / *mms6*-Halo BPC *n* = 1199 cells, Δ*mamM* / *mms6*-Halo NBPC *n* = 285 cells, Δ*mamM* / *mms6*-Halo BPC *n* = 401 cells, Δ*mamE* Δ*limE* / *mms6*-Halo NBPC *n* = 154 cells, Δ*mamE* Δ*limE* / *mms6*-Halo BPC *n* = 383 cells, Δ*mamA* / *mms6*-Halo NBPC *n* = 518 cells, Δ*mamA* / *mms6*-Halo BPC *n* = 314 cells, Δ*amb0955* Δ*mms6* Δ*mmsF* / *mms6*-Halo NBPC *n* = 278 cells, Δ*amb0955* Δ*mms6* Δ*mmsF* / *mms6*-Halo BPC *n* = 530 cells.

To identify MAI proteins involved in Mms6 translocation to magnetosome membranes, Mms6-Halo was expressed in strains deleted for specific MAI genes. Given Mms6 only localizes to magnetosomes that contain magnetite (16), we first focused on four strains in which magnetite synthesis is completely disrupted, Δ*mamO*, Δ*mamM*, Δ*mamE, and* Δ*mamN* (28). As expected, when *mamO*, *mamM*, or *mamE* is deleted, Mms6-Halo only appears in the cytosol, suggesting that either Mms6 is not localizing to magnetosomes because they lack a mineral or that MamM, MamO, and MamE are more directly involved in Mms6 localization (Fig. 5A,B). Unexpectedly, when another protein essential for magnetite synthesis, MamN, is absent, Mms6-Halo localizes to magnetosome membranes regardless of biomineralization conditions and despite the absence of crystal production (Fig. 5A,B). This surprising exception may indicate that MamN inhibits Mms6 localization until biomineralization begins. Additionally, it demonstrates that the presence of magnetite is not necessary for Mms6 localization.

A previous study by Nguyen et al. found that Mms6 interacts with MamA (31). Therefore, we also examined the localization of Mms6-Halo in a *mamA* deletion mutant. Mms6-Halo localized similarly in the Δ*mamA* and WT strains under NBPC. While there is a significant difference between WT and Δ*mamA* cells in the categorical distribution of Mms6-Halo localization under BPC, the effect size is small and the majority of Δ*mamA* cells still show Mms6-Halo aligned to magnetosome chains. These data taken together indicate that *mamA* is not strictly required for Mms6 localization to magnetosomes (Fig. 5A,B). However, it is still possible that MamA is recruited by Mms6, or that the two proteins interact for a purpose other than localization. Due to its genomic proximity to *mms6*, *mmsF* was also tested. We examined the localization of Mms6-Halo in a strain lacking the *mms6* gene cluster, containing *mms6, mmsF,* and the uncharacterized protein *amb0955.* Under NBPC, significantly more *mms6* cluster mutant cells have Mms6-Halo localized to the magnetosome chain, but most cells still have diffuse Mms6-Halo (Fig. 5A,B). Under BPC, Mms6-Halo localizes normally, suggesting that other *mms6* cluster proteins are not needed for *mms6* localization to magnetosomes, but could have a small positive effect on Mms6 cytosolic localization. Additionally, it is possible that *mmsF* homologs *amb0953* and/or *amb1013* could serve a redundant function for *mmsF* and mask the effect of its deletion (32). Examples of localization patterns in the above backgrounds imaged with super resolution microscopy are provided in Supplementary Figure 2.

### Defining intrinsic determinants of Mms6 localization

After identifying other MAI proteins that affect Mms6 localization, we looked for Mms6 domains that contribute to localization. Mms6 can be roughly divided into four protein domains: the N-terminal domain (NTD), glycine-leucine repeat segment (GL), transmembrane domain (TM), and the magnetite-interacting component (MIC) (Fig. 6A). The N-terminal domain is a 98 amino acid region that was not identified when Mms6 was originally discovered in proteomic analyses of magnetite-associated peptides (17). Thus, the NTD may be cleaved from Mms6 during or after localization to the magnetosome membrane. The GL repeat domain consists primarily of alternating glycine and leucine residues and is a defining feature of silk fibroin that may mediate protein-protein interactions (33). An approximately 23 amino acid transmembrane domain (TM) is predicted to begin in the middle of the GL repeat domain (13). Consistent with the presence of a transmembrane region, Mms6 has been identified in enriched magnetosome membranes (13). The MIC is a region of acidic amino acids that binds ferrous iron (18,34), ferric iron (17,35,36), magnetite crystal (37,38), and other minerals (17). The MIC has been implicated in iron crystal nucleation (17,18) and protein localization of Mms6 (23). To test the effect of Mms6 domains on its dynamic localization, truncated Mms6 proteins were expressed in a Δ*mms6* background. Because Mms6 only localizes to magnetosomes that contain magnetite, it seemed likely that the MIC would contribute to localization (16). Unexpectedly, the MIC was dispensable for normal Cmag (Fig. 6B). Under NBPC, Mms6_1-139_-GFP, lacking the MIC, localizes to the cytoplasm like Mms6-GFP. Surprisingly, in the majority of cells Mms6_1-139_-GFP localizes to magnetosomes under BPC (Fig. 6C,D), except for a small but significant increase in diffuse localization, suggesting that magnetite-binding activity is not necessary for Mms6 localization.

**Figure 6.**
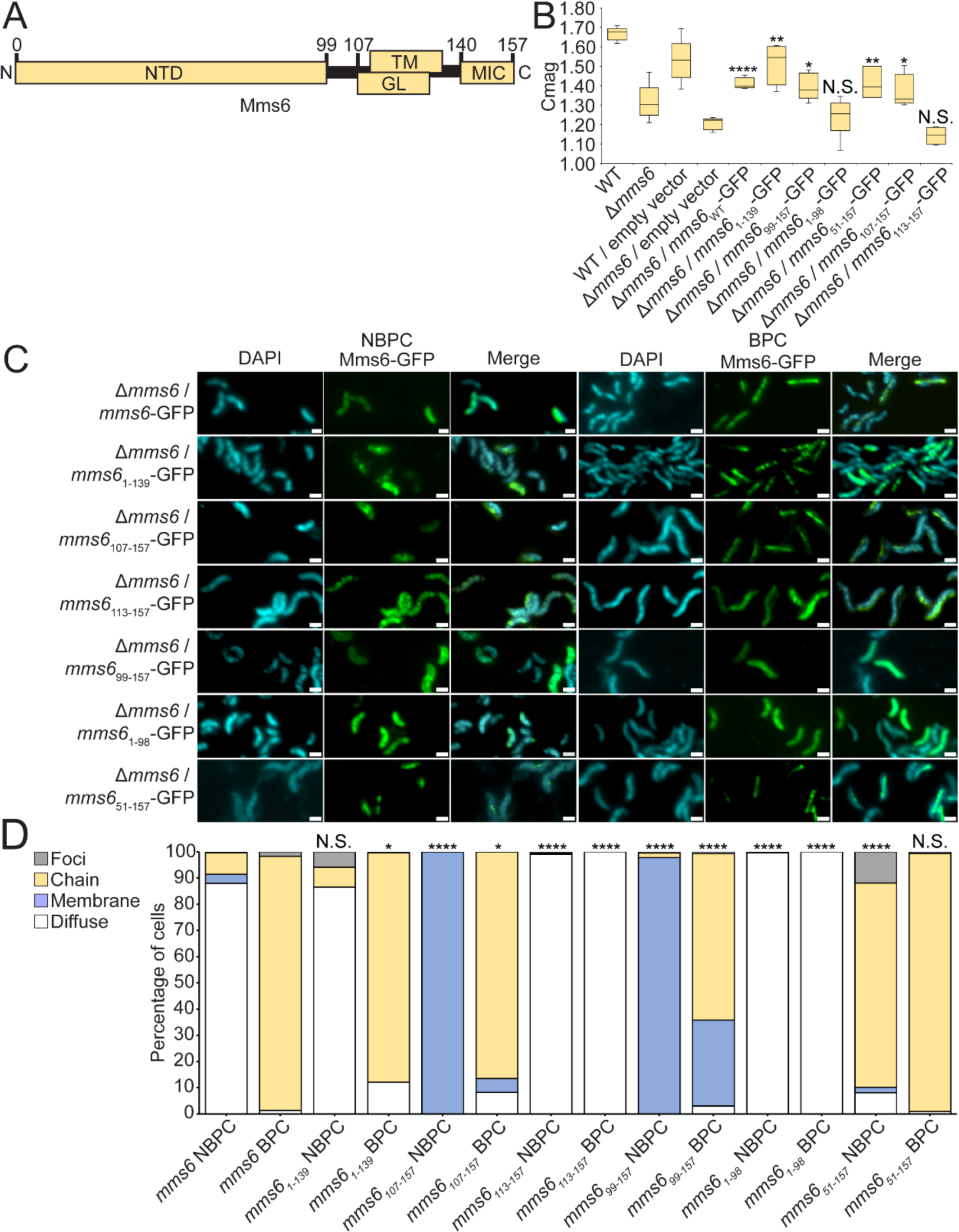
Mms6 protein domains are essential to conditional and diffuse localization. (A) Diagram of Mms6 protein domains (NTD N-terminal domain, TM transmembrane domain, GL glycine-leucine repeat region, MIC magnetite-interacting component). (B) Coefficient of magnetism (Cmag) of several strains measured by differential scattering of light by cells moved into different orientations by an external magnetic field. *P* values were calculated by Mann-Whitney U Test comparing given dataset to Δ*mms6* / empty vector (N.S. no significant difference *P* > .01) (* = *P* < 10^-2^) (** = *P* < 10^-3^) (**** = *P* < 10^-5^). (C) Representative fluorescence microscopy images of *Δmms6* expressing WT *mms6* or a mutant version. DAPI counterstain is shown in blue. Scale bars = 1 µm. (D) Blind quantification of localization patterns of WT and mutant versions of Mms6-GFP in *Δmms6. P* values were calculated by Fisher’s exact test comparing Mms6 mutant datasets to Δ*mms6* / *mms6*-GFP (N.S. no significant difference *P* > .01) (* = *P* < 10^-2^) (**** = *P* < 10^-5^). Δ*mms6* / *mms6*-GFP NBPC *n* = 233 cells, Δ*mms6* / *mms6*-GFP BPC *n* = 812 cells, Δ*mms6* / *mms6_1-139_*-GFP NBPC *n* = 128 cells, Δ*mms6* / *mms6_1-139_* BPC *n* = 470 cells, Δ*mms6* / *mms6*_99-157_-GFP NBPC *n* = 411 cells, Δ*mms6* / *mms6*_99-157_-GFP BPC *n* = 265 cells, Δ*mms6* / *mms6*_1-98_-GFP NBPC *n* = 437 cells, Δ*mms6* / *mms6*_1-98_-GFP BPC *n* = 1112 cells, Δ*mms6* / *mms6*_51-157_-GFP NBPC *n* = 285 cells, Δ*mms6* / *mms6*_51-157_-GFP BPC *n* = 746 cells, Δ*mms6* / *mms6*_107-157_-GFP NBPC *n* = 576 cells, Δ*mms6* / *mms6*_107-157_-GFP BPC *n* = 1095 cells, Δ*mms6* / *mms6*_113-157_-GFP NBPC *n* = 1152 cells, Δ*mms6* / *mms6*_113-157_-GFP BPC *n* = 1189 cells.

After finding that the MIC is dispensable for magnetosome localization, we made further truncations of Mms6. One truncation, Mms6_107-157_-GFP, consists of the GL repeat region, the transmembrane region, and the MIC. Under BPC, this variant localizes to magnetosome membranes in most cells, with small but significant increases in diffuse and membrane localization compared to full length Mms6-GFP. Under NBPC, where the full length Mms6-GFP is cytoplasmic, Mms6_107-157_-GFP localizes to the cellular membrane (Fig. 6C,D) (Supplementary Figure 5). In contrast, Mms6_113-157_-GFP, which consists of the TM region and MIC, is diffuse in all conditions (Fig. 6C,D), suggesting it is unable to translocate to the membrane or localize to magnetosomes. These results indicate that a factor within the GL repeat region may be necessary for Mms6 membrane localization. A mutant Mms6_107-135_-GFP was made to test if Mms6 localization was possible with only the GL and TM domains, but no GFP signal was seen, likely due to protein instability or loss through proteolysis.

Next, we investigated a segment of Mms6 thought to be cleaved from the mature protein. Mms6 was originally discovered in an experiment by Arakaki et al. that dissolved the magnetite crystal and analyzed proteins in the resulting solution. The solution contained a 6 kDa peptide of Mms6, but the gene codes for a larger protein of 12-15kDa (11,17). Both the 6 kDa and 14.5 kDa Mms6 proteins exist in the cell (10). The shorter form of the protein lacks the 99 amino acid N-terminal domain (NTD), which may be cleaved from the mature protein by MamE protease activity (17,30). Previous work by Arakaki et al. found that without the NTD, Mms6 localizes diffusely in either the cytoplasm or cellular membrane (16). To investigate the effect of the NTD, several mutants of Mms6 were examined for conditional localization. Under NBPC, Mms6_99-157_-GFP appears in the cell membrane instead of the cytosol (Fig. 6C,D). Under BPC, Mms6_99-157_-GFP localizes both to magnetosome chains and to the cell membrane. These localization differences may indicate that the NTD keeps Mms6 diffuse in the cytosol, possibly to prevent it from translocating into membranes before the initiation of magnetite biomineralization. When Mms6_99-157_-Halo is expressed in ΔMAI mutants it localizes to the cellular membrane (Supplementary Fig. 6). This result suggests that Mms6_99-157_, unlike Mms6, does not require other magnetosome proteins or pre-formed magnetosome membranes to associate with membranes. Mms6_1-98_-GFP, containing only the N-terminal domain, localizes to the cytosol regardless of biomineralization condition (Fig. 6C,D). To further investigate the NTD, we created Mms6_51-157_-GFP, in which the N-terminal half of the NTD is absent. Interestingly, Mms6_51-157_-GFP localized to magnetosome membranes under both BPC and NBPC (Fig. 6C,D), suggesting that the NTD may also be involved in the conditional localization of Mms6. Super resolution images of example cells expressing the Mms6 mutants discussed above are shown in Supplementary Figure 6.

To test the effect of *mms6* domain fusions with other magnetosome proteins, the N-terminal domain was fused to the N-terminus of *mmsF.* Wild-type MmsF tagged N-terminally with GFP localizes to magnetosome chains regardless of biomineralization conditions (Fig. 7). However, the addition of the Mms6 N-terminal domain imparts conditional localization to MmsF. Under NBPC, GFP-Mms6_NTD_MmsF is found at the cellular membrane and, similar to Mms6, shows magnetosome localization only in BPC. Therefore, the Mms6 N-terminal domain is both necessary and sufficient for conditional localization in magnetosome proteins. Interestingly, under NBPC, the fusion protein appears localized to the cell inner membrane, suggesting that the Mms6 NTD prevents magnetosome localization of MmsF but not its membrane translocation. This may indicate that the Mms6 NTD has two separate functions - keeping proteins cytosolic, and controlling conditional localization - and that perhaps the NTD is only able to maintain Mms6 in a cytosolic location in concert with other structural features of Mms6. These results open future possibilities for modifying magnetosome protein localization using the NTD.

**Figure 7.**
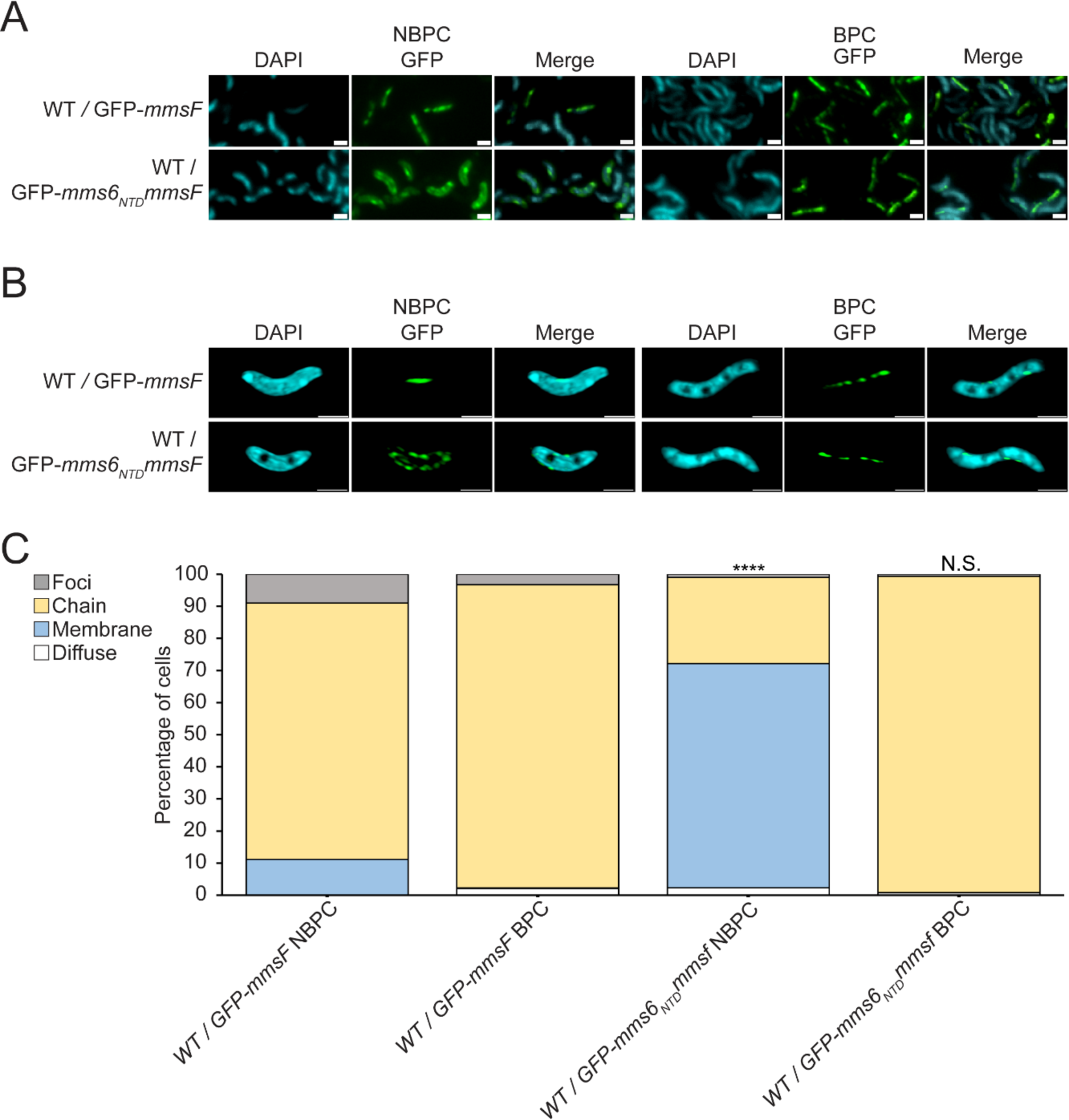
*mms6* N-terminal domain fused to *mmsF* imparts conditional localization. (A) Representative fluorescence microscopy images of WT cells expressing either GFP-*mmsF* or GFP-*mms6_NTD_mmsF* shown in green and DAPI shown in blue. (B) Representative super resolution 3D Structured Illumination Microscopy (SIM) images of WT cells expressing either WT / GFP-*mmsF* or WT / GFP-*mms6_NTD_mmsF* shown in green and DAPI shown in blue. Scale bars = 1 µm. (C) Blind quantification of localization patterns based on fluorescence microscopy images. *P* values were calculated by Fisher’s exact test comparing WT / GFP-*mms6_NTD_mmsF* to WT / GFP-*mmsF* dataset of respective biomineralization condition (N.S. no significant difference *P* > .01) (**** *P* < 10^-5^). WT / GFP-*mmsF* NBPC n = 348 cells, WT / GFP-*mmsF* BPC n = 802 cells, WT / GFP-*mms6_NTD_mmsF* NBPC n = 906 cells, WT / GFP-*mms6_NTD_mmsF* BPC n = 625 cells.

### Biochemical fractionation of AMB-1 to determine Mms6 localization

Due to the contrast between the existence of a transmembrane domain in Mms6 and its cytosolic location under NBPC, we sought to validate microscopic observations of Mms6 using biochemical subcellular fractionation. Briefly, AMB-1 cells were lysed, and ultracentrifugation was used to separate soluble and insoluble cellular contents. Known magnetosome membrane protein MamE was used as an insoluble fraction marker (10). Based on the cytosolic pattern of Mms6-Halo in most cells under NBPC, it was expected that Mms6-Halo would appear in the soluble fraction. Surprisingly, Mms6-Halo was only found in the insoluble fraction (Supplementary Fig. 7).

A variety of factors could cause Mms6-Halo to co-fractionate with insoluble proteins, despite having a cytoplasmic location. Mms6 phase separates *in vitro* and forms protein micelles (35,39). Therefore, we attempted to prevent the formation of protein micelles using the mild non-ionic detergent Igepal CA-630 (NP-40 substitute). Cellular fractionation performed with Igepal resulted in the solubilization of Mms6-Halo, whereas magnetosome membrane protein MamE was still present primarily in the insoluble fraction (Supplementary Fig. 7). However, Mms6-Halo was also soluble in cells grown in BPC where it was expected to be inside the magnetosome lumen (Supplementary Fig. 7). The greater solubility of Mms6 compared to MamE may suggest that Mms6 is associated less strongly with magnetosome membranes or that Mms6 is only surface associated.

### MamD localizes conditionally to magnetosomes similar to Mms6

Many MAI proteins have been implicated in growing and shaping developing magnetite crystals including the proteins of the *mamGFDC* operon and *mms6* gene cluster. Past studies have demonstrated that MamF, MamC, Mms6, and MmsF all localize to the magnetosome when examined with fluorescence microscopy in AMB-1 (16,32,40). MamD co-fractionates with the magnetosome fraction biochemically, but this result has not yet been corroborated with fluorescence microscopy (17,23). MamG localization in AMB-1 has not been experimentally determined, but it is proposed to localize to magnetosomes similar to its homolog MamG in MSR-1 (12). Thus, we asked if any other crystal shaping proteins exhibit conditional localization to magnetosomes in a manner similar to Mms6.

To investigate protein sorting, WT cells expressing GFP-tagged proteins were grown in either BPC or NBPC. Mms6-GFP, as described above, was mostly chain aligned under BPC and cytosolic under NBPC (Fig. 8A,B). GFP-MmsF, in contrast, localizes to magnetosomes regardless of biomineralization conditions (Supplementary Fig. 8). MamG-GFP, MamF-GFP, and MamC-GFP also displayed magnetosome localization in most cells in both conditions (Supplementary Fig. 9). Although the distribution of protein localization patterns was different between BPC and NBPC for these proteins, the effect sizes were small, and the majority of cells show chain-aligned protein. This suggests that the proteins localize to magnetosomes under both BPC and NBPC.

**Figure 8.**
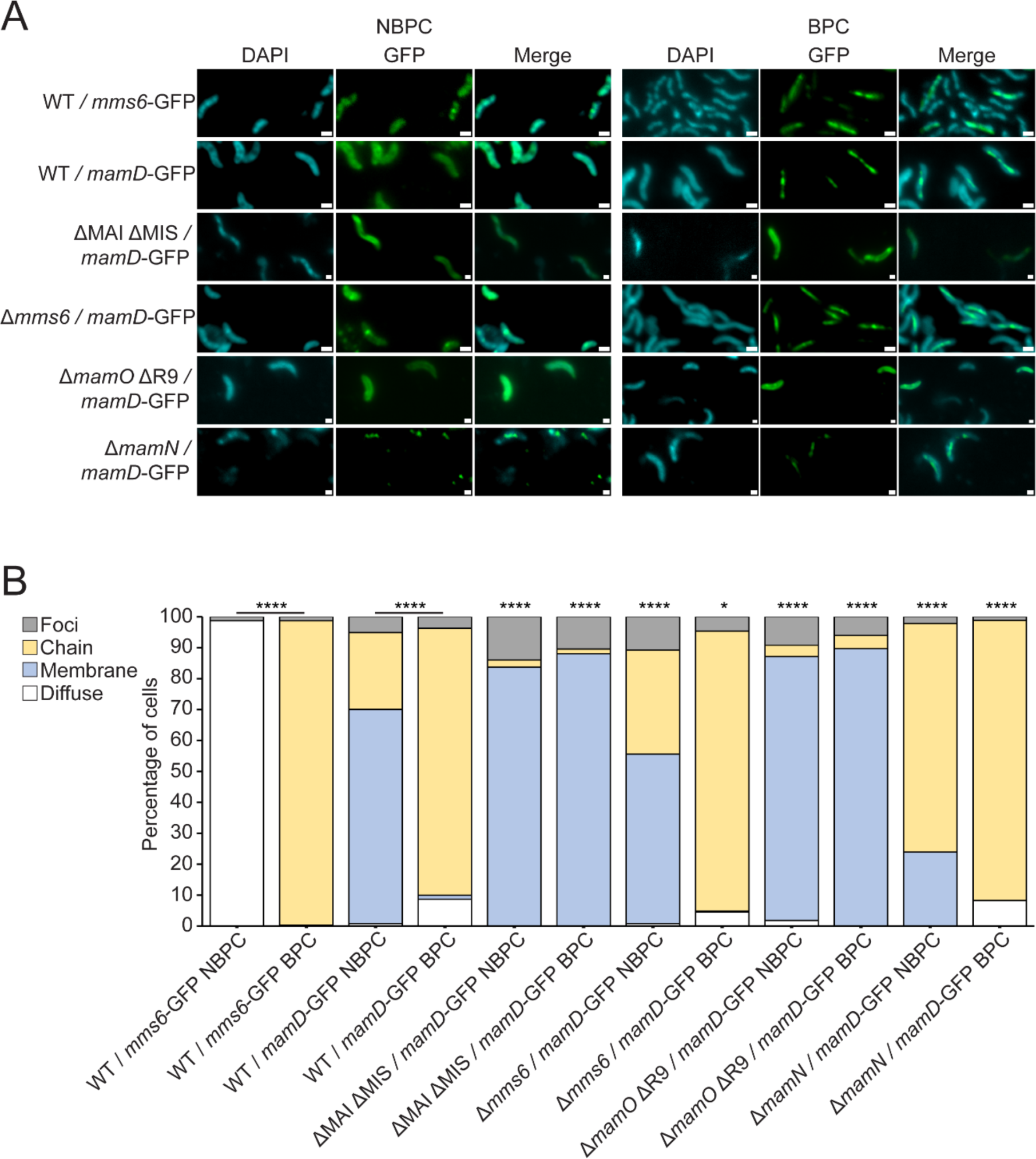
MamD-GFP localizes to magnetosomes conditionally similar to Mms6. (A) Representative fluorescence microscopy images of WT or mutant AMB-1 cells grown under standard growth conditions expressing MamD or Mms6 GFP fusions. GFP is shown in green, and DAPI is shown in blue. Scale bars = 1 µm. (C) Blind quantification of localization patterns of GFP tagged Mms6 or MamD expressed *in vivo*. Cells were categorized by localization pattern. The Y-axis represents percentage of total cell count with indicated protein fluorescence pattern. *P* values were calculated by Fisher’s exact test comparing indicated datasets (* *P* < .01) (**** *P* < 10^-5^). Localization patterns of MamD-GFP expressed in mutant cells were compared statistically to WT under the same biomineralization conditions. Effect sizes are listed in supplementary table S4. WT / *mms6*-GFP NBPC *n* = 1074 cells, WT / *mms6*-GFP BPC *n* = 1317 cells, WT / *mamD*-GFP NBPC *n* = 1041 cells, WT / *mamD*-GFP BPC *n* = 971 cells, ΔMAI ΔMIS / *mamD*-GFP NBPC *n* = 43 cells, ΔMAI ΔMIS / *mamD*-GFP BPC *n* = 67 cells, *Δmms6* / *mamD*-GFP NBPC *n* = 1372 cells, *Δmms6* / *mamD*-GFP BPC *n* = 1660 cells, Δ*mamO* ΔR9 / *mamD*-GFP NBPC *n* = 218 cells, Δ*mamO* ΔR9 / *mamD*-GFP BPC *n* = 165 cells, Δ*mamN* / *mamD*-GFP NBPC *n* = 2599 cells, Δ*mamN* / *mamD*-GFP BPC *n* = 1833 cells.

In contrast to these proteins, MamD displayed conditional localization similar to Mms6 (Fig. 8B,C). MamD-GFP localizes to magnetosomes in most cells only under biomineralization conditions and is distributed on the cell membrane under non-biomineralization conditions. Notably, MamD-GFP was membrane-localized under non-biomineralization conditions, whereas Mms6-GFP appears diffuse in the cytoplasm. This may suggest that the localization of MamD is regulated differently from that of Mms6.

We envision two potential cellular routes for the conditional localization of MamD-GFP to magnetosomes. First, MamD may be recruited directly by Mms6, which itself displays conditional localization. Past work by Tanaka et al. found that MamC and MamD were depleted from the fraction tightly bound to magnetite crystal in Δ*mms6* (18), suggesting that Mms6 may recruit crystal-shaping proteins to the magnetosome. Second, like Mms6, MamD may be sorted to the magnetosome only during biomineralization via the MamEOMN proteins.

To differentiate between these possibilities, we examined the localization of MamD-GFP expressed in WT and in mutants lacking *mms6* or other genes found to be important for Mms6 localization. MamD-GFP does not require *mms6* for magnetosome localization (Fig. 8A,B) suggesting that it is not recruited to magnetosomes by Mms6. However, in ΔMAI ΔMIS or Δ*mamO*, MamD-GFP was dispersed on the inner membrane as it was in the absence of magnetite formation (Fig. 8A,B). In contrast, in a Δ*mamN* background, most cells have MamD-GFP at magnetosomes regardless of biomineralization conditions. This localization pattern is reminiscent of Mms6-Halo expressed in a Δ*mamN* background, suggesting that the magnetosome localization of MamD, like Mms6, is inhibited by MamN (Fig. 8A,B). These findings suggest that there are at least two magnetite-maturation protein sorting systems in AMB-1, one that sorts Mms6 and MamD based on biomineralization conditions through inhibition by MamN, and a second in which proteins like MamG, MamF, and MamC are sorted to magnetosomes before biomineralization begins. Together, our results reveal the complexity of magnetosome protein sorting as well as raise new questions about magnetosome protein modification and membrane topology.

## Discussion

The identity and function of magnetosome organelles is dependent on the activity of the collection of proteins localized to the compartment. A generally accepted model proposes that proteins localize to magnetosomes during the formation of the organelle regardless of environmental conditions. However, a previous study showed that the magnetite-shaping protein, Mms6, localizes to magnetosomes only under cellular conditions that promote biomineralization (16). Here, we further refine the intrinsic and extrinsic parameters that define this unusual mode of protein localization.

### An alternate route for protein localization to magnetosomes

Using a pulse-chase experiment, we show that a pool of Mms6 produced under NBPC can relocalize to full magnetosome chains within 1-2 h after a switch to BPC. Several observations indicate that the localization of Mms6 does not require the formation of new magnetosomes. First, AMB-1 has a doubling time of 4-6 h, significantly longer than the time for Mms6 relocalization in our experiments (Figure 1). Second, previous work by Cornejo et al., using a synthetic inducible magnetosome formation system, found that a new magnetosome chain is constructed in approximately 3-6 hours (41). Based on this timescale, it is unlikely that a complete chain of new magnetosomes could be formed in the 2 h needed for Mms6 to relocalize. Therefore, we favor a scenario in which the protein is translocated into pre-existing magnetosomes. If localization was restricted to newly formed magnetosomes within an existing chain, we would expect an intermediate phase with only a few foci of fluorescence within the cell. The absence of this step suggests the Mms6 pool can relocalize to all crystal-containing magnetosomes in the chain at once. These results taken together indicate that cytoplasmic Mms6 can relocalize to pre-existing magnetosomes when biomineralization conditions change, revealing a surprising mode of magnetosome transmembrane protein localization in AMB-1.

### Mms6 domains required for regulated magnetosome localization

To investigate the intrinsic determinants of localization, we created several variants of Mms6 lacking its previously characterized domains. We show that the C-terminal magnetite-interacting component (MIC) of Mms6 is not necessary for localization to the magnetosome or to the cytoplasm (Fig. 6C,D). This finding, along with the observations of the *ΔmamN* strain, further demonstrate that magnetite-binding is not needed for localization of Mms6 to magnetosomes. The MIC is also dispensable for normal magnetic response (Fig. 6B) suggesting that, independent of mineral-binding, Mms6 may have other functions that affect the magnetic response. It is also possible that other Mms proteins may substitute for the lack of magnetite-binding in this mutant.

A study by Yamagishi et al. showed that deletions in the Mms6 MIC resulted in cells with misshapen magnetite crystals (23), suggesting that this domain is needed for Mms6 localization, function, or stability. To test whether localization was affected by these mutations, Yamagishi et al. isolated the magnetosome membrane fraction from the cytoplasm and the inner membrane fraction and performed immunoblotting. Wild-type Mms6-His was found specifically in the magnetosome membrane, whereas His-tagged mutants with deletions in the mineral interacting component were absent from the magnetosome membrane. These results were taken to mean that Mms6 requires the MIC for magnetosome localization. In contrast, our live cell fluorescence microscopy results show that Mms6 variants lacking the MIC localize to magnetosomes (Fig. 6C). In addition, complementing the *mms6* deletion mutant with Mms6_1-139_-GFP restores the cellular magnetic response (Fig. 6B). This discrepancy may be due to a difference in methodology. Yamagishi et al. found that the Mms6 MIC mutants were also absent from the other cell fractions, suggesting that they may have been unstable or degraded by proteases. Mutant Mms6_107-135_-GFP created for our study expressed a similar length of Mms6 as the mutant from Yamagishi et al. and gave a signal too faint to image, likely due to protein degradation or instability. Thus, the N- and C-terminal ends of Mms6 may stabilize the protein *in vivo*. Further work will be necessary to determine the minimal protein domains necessary for the magnetosome sorting of Mms6.

A previous study by Arakaki et al. showed that without the NTD, Mms6-GFP localizes diffusely under BPC (16). However, it could not be determined whether Mms6-GFP localized in either the cytoplasm or cell membrane (16). Here, we show that under NBPC, Mms6_99-157_-GFP localizes to the cell membrane instead of the cytosol (Fig. 6C,D). In contrast to previous results, we find that under BPC Mms6_99-157_-GFP localizes to the cell membrane and to magnetosome chains. This discrepancy could be due to the difficulty at lower resolution in distinguishing proteins aligned to magnetosome chains from proteins aligned to the cellular membrane. We further demonstrate that the N-terminal half of the Mms6 NTD is necessary for its conditional localization. A truncated Mms6 lacking the NTD produced *in vitro* has been shown to form large micellar homopolymers (35,42). It is unclear if these micelles form *in vivo* or if they relocalize with changing biomineralization conditions. Therefore, the NTD may serve to keep Mms6 monomers free within the cytosol for rapid re-sorting when required. Notably, in some species, such as *Magnetovibrio blakemorei*, Mms6 lacks the NTD, suggesting that it is not necessary for effective biomineralization (43). Surprisingly, the localization pattern of MmsF becomes conditional when fused with the Mms6 NTD, suggesting the NTD could be used to direct heterologous proteins to magnetosome membranes under specific conditions. However, the Mms6 NTD-MmsF fusion protein does not become cytoplasmic in NBPC like Mms6, indicating that other properties of Mms6 mediate its retention in the cytoplasm. Alternatively, the native membrane localization properties of MmsF may override the cytoplasmic retention activity of the NTD.

### Alternative models of Mms6 topology

Our work raises broader questions regarding the topology and localization of Mms6. Mms6 was isolated as a magnetite interacting protein, suggesting that parts of the protein, namely the MIC, face the interior of the magnetosome. It is also predicted to have a transmembrane domain and may interact with MamA, which is on the cytoplasmic side of the membrane. However, we show that folded Mms6, tagged with either GFP or Halo, can localize to magnetosomes during a switch from NBPC to BPC. These tags are at the C-terminus of Mms6 directly following its MIC. Since folded proteins generally cannot use the Sec translocon to cross membranes, we suggest that an alternate pathway is used for the regulated localization of Mms6 to membranes. The translocation of fully folded proteins is poorly understood in bacteria outside of the twin-arginine transport (TAT) systems (44). Mms6 lacks a TAT signal peptide, indicating it is unlikely to be transported by the TAT system. Mms6 transport also does not follow the pattern of other known bacterial folded protein transporters such as the Type 3 secretion system (T3SS) and Type 9 secretion system (T9SS), both of which translocate proteins from the cytoplasm across both the inner and outer membranes into the extracellular space (44). These findings suggest that Mms6 may localize through an undiscovered membrane transporter that translocates fully folded proteins.

The biochemical analysis of Mms6 localization also raises questions about its topology and biophysical state in the cell. Under BPC, Mms6 co-fractionates with membrane proteins such as MamE, as would be expected from the current models of its localization to magnetosomes as a membrane-bounded protein. Surprisingly, however, under NBPC Mms6 appears diffuse in the cytoplasm by microscopy but still co-fractionates with MamE in the insoluble fraction (Supplementary Fig. 7). Mild detergent treatment turns Mms6 into a soluble protein under all conditions while MamE remains within the insoluble fraction. These findings suggest that Mms6 may be present in micellar form or only have a weak association with the magnetosome membrane. These observations, along with its translocation as a folded protein, raise the possibility that Mms6 is not a transmembrane-domain-containing protein. Perhaps, Mms6 remains in micellar form and translocates to the lumen of the magnetosome where it participates in biomineralization through direct interactions with magnetite. Alternatively, Mms6 may only associate with the cytoplasmic side of the magnetosome membrane. In this light, the previous association of Mms6 with magnetite may have been an artifact of its iron-binding properties. Further research will need to be done to clarify Mms6 topology.

### Membrane growth and protein localization

The conditional localization of Mms6 requires several genes previously implicated in magnetite synthesis and magnetosome membrane growth. In the absence of *mamE, mamM,* and *mamO,* Mms6 fails to localize to magnetosomes. In contrast, in the absence of the magnetite synthesis gene *mamN,* Mms6 localizes to magnetosomes under all conditions. Additionally, MamD, another magnetite-shaping protein, requires MamO for its conditional localization to magnetosomes and is prevented from magnetosome entry by MamN under NBPC. The MAI proteins that impact the sorting of Mms6 and MamD are known components of a checkpoint that regulates the growth of magnetosome membranes in AMB-1 (41,45,46). Prior to biomineralization, magnetosome membranes grow to a size of approximately ∼35 nm. A second stage of magnetosome membrane growth occurs after the initiation of biomineralization. This second stage of growth requires the activation of MamE protease activity by MamO. Active MamE proteolyzes MamD, and other substrates to allow for expansion of the magnetosome membrane. MamN is a negative regulator of membrane growth via an unknown mechanism (30). In its absence, empty magnetosome membranes can escape the checkpoint and grow larger. Thus, factors that positively regulate membrane growth (BPC; MamE and MamO) also promote the conditional localization of Mms6. Factors that negatively regulate membrane growth (NBPC; MamN) also inhibit conditional localization of Mms6. Importantly, the membrane growth checkpoint does not depend on the presence of Mms6 (30).

Based on these observations we present an integrated model of membrane growth and protein localization in Figure 9. In this model, MmsF and other crystal maturation proteins like MamG and MamF are first recruited to the magnetosome, during or after magnetosome membrane formation (Fig. 9). MamN inhibits the magnetosome localization of Mms6 and MamD until biomineralization permissible conditions are reached. MamN inhibition is then lifted, possibly through the protease activity of MamE after activation by MamM and MamO, and Mms6 and MamD can localize to magnetosomes. Alternatively, MamE, MamM, and MamO may directly recruit Mms6 and MamD to magnetosomes. MamD localized to magnetosomes keeps membranes under the size threshold to enhance the concentration of iron in the lumen and promote efficient magnetite nucleation. Once nucleation has begun, MamE cleaves MamD allowing a second stage of membrane growth.

**Figure 9.**
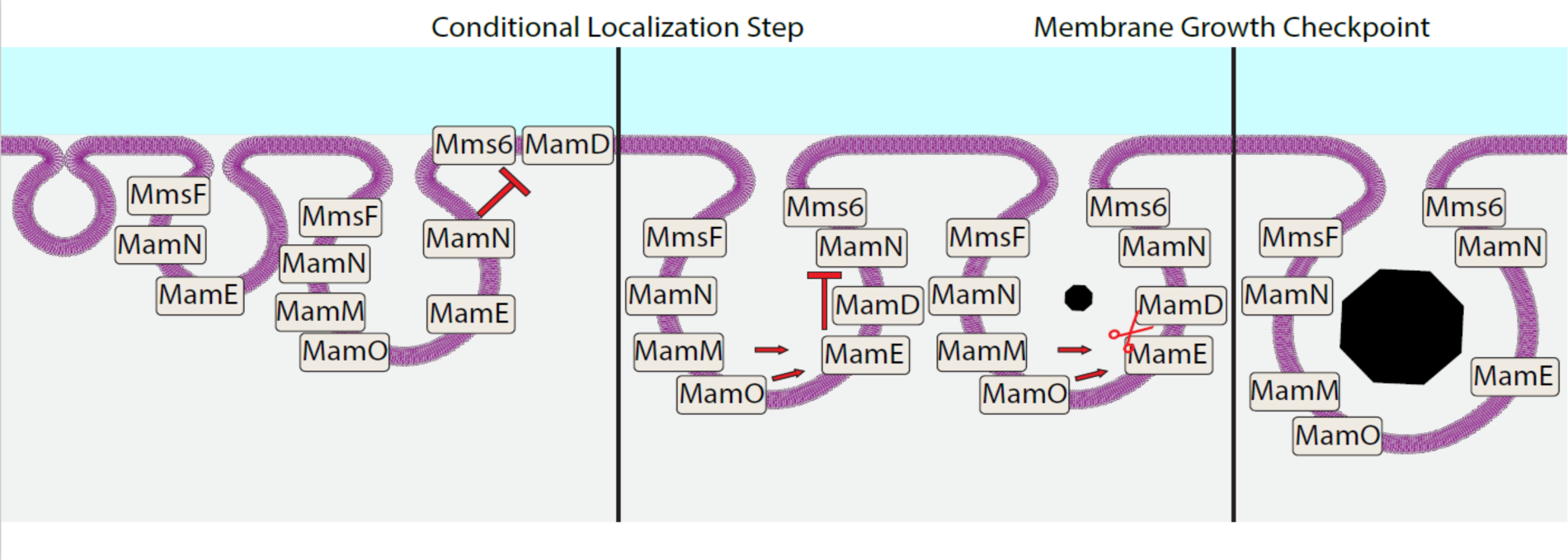
Model for sorting of magnetite maturation proteins. MamN, MamO, and likely other proteins promote the recruitment of MmsF through an unknown mechanism. After magnetosome membrane formation, but before biomineralization begins, MamN inhibits localization of Mms6 and MamD. Once biomineralization conditions are reached, MamN is inhibited by the action of another factor, possibly MamE, which could be activated by MamO and MamM. MamN inactivation allows Mms6 and MamD to localize to the magnetosome, where they aid in magnetite nucleation. MamD works to restrict membranes under a size threshold to concentrate mineral, facilitating nucleation. Once magnetite is nucleated, MamE protease domain is activated by MamO and MamM, cleaving MamD and allowing further membrane expansion.

Our findings emphasize the importance of tight cellular control over protein localization in biomineralization. We show that at least two MAI proteins are dynamically sorted to magnetosome compartments as biomineralization conditions change. Previous work has shown that another magnetosome protein in AMB-1, McaB, also has a similar conditional localization (47). Therefore, conditional localization may be a more common mode of protein sorting to magnetosomes. We also demonstrate that the localization of biomineralization proteins can be modified *in vivo* using the Mms6 NTD. Developing further abilities to modify magnetosome protein localization and target new proteins to the magnetosome membrane could allow finer control over the production of magnetite particles that can be used in medical and biotechnological applications.

## Materials and Methods

### Bacterial growth and cellular magnetic response

The strains used in this study are described in Table S1. AMB-1 stock cultures were grown as described by picking single colonies and growing in 1.5 mL Magnetospirillum Growth (MG) medium in 1.7 mL microtubes (Genesee Scientific Cat #24-281) at 30 °C for 3-4 d with 15 µl Wolfe’s vitamin solution and 20 µM ferric malate (48). To start larger cultures, stock cultures were diluted 1:100 in 10 mL MG medium with 100 µl Wolfe’s vitamin solution and 20 µM ferric malate in 24 mL capped tubes and grown at 30 °C for 2 d in a 10% oxygen microaerobic chamber. Antibiotic selection was done with 10 µg/mL kanamycin in solid MG medium and 7 µg/mL kanamycin in liquid MG medium.

To record the magnetic response (Cmag) of an AMB-1 culture, the optical density of AMB-1 cells growing in 10 mL MG medium was measured in a UV-vis spectrophotometer. An external magnetic field was applied to the cells to shift magnetic cells from a parallel to perpendicular orientation relative to the light beam, creating a quantifiable difference in optical density used to represent magnetic response. The ratio of measured OD values when the magnetic field is parallel versus perpendicular is recorded.

*Escherichia coli* cultures were grown in 10 mL lysogeny broth in 24 mL capped tubes on a rotating wheel at 37 °C for about 12-16 h. Antibiotic selection was done with 50 µg/mL kanamycin. An addition of 300 µM diaminopimelic acid was necessary to grow *E. coli* strain WM3064.

### Genetic manipulation

Oligonucleotides were designed in sequence analysis software Geneious using the *Magnetospirillum magneticum* AMB-1 genome sequence NC_007626.1 and were manufactured by Elim Biopharm or Integrated DNA Technologies. DNA fragments were amplified using GoTaq master mix (Promega Cat #M7123). Plasmids were introduced into AMB-1 through conjugation and are listed in supplementary table S2.

Several plasmids were created to express truncated versions of Mms6 with a GFP fusion tag. To create pAK1456 (*mms6*_99-157_-GFP), pAK1444 (*mms6*_1-98_-GFP), pAK1445 (*mms6*_113-157_-GFP), pAK1446 (*mms6*_107-157_-GFP), pAK1441 (*mms6*_51-157_-GFP), and pAK1443 (*mms6*_1-139_-GFP), fragments of mms6 were PCR amplified from AMB-1 genomic DNA using the primers listed in supplementary table S3 and inserted by Gibson assembly into pAK1102 (*mms6*-GFP), following vector digestion with BamHI-HF and EcoRI-HF restriction enzymes (New England Biolabs). To create pAK1447 (GFP-*mms6*_NTD_*mmsF*), fragments of mms6 were PCR amplified from AMB-1 genomic DNA using the primers listed in supplementary table S3 and inserted by Gibson assembly into pAK532 (GFP-*mmsF*), following vector digestion with BamHI-HF and SpeI-HF restriction enzymes (New England Biolabs). To create pAK1440 (*mamG*-GFP), *mamG* was amplified from AMB-1 genomic DNA using the primers listed in supplementary table S3 and inserted by Gibson assembly into multiple cloning vector pAK22, following vector digestion with BamHI-HF and EcoRI-HF restriction enzymes (New England Biolabs).

### Fluorescence microscopy and localization pattern quantification

To analyze Mms6 localization in AMB-1 cells, cells growing in 10 mL MG medium were collected once reaching an optical density of OD_400_ 0.08-0.15 by centrifugation at 10,000 x g for 3 min. Optical density is measured on a Thermo Spectronic 20D+. Cell pellets expressing a HaloTag fusion were resuspended in 100 µL MG medium, incubated with 500 nM HaloTag 549 ligand (Promega Cat #GA1110) for 60 min in the dark at 30 °C in a 10% O_2_ microaerobic chamber, and then centrifuged at 10,000 x g for 3 min. All cell pellets were then resuspended in 100 µL fresh MG medium and stained with 1.4 µM 4’,6-diamidino-2-phenylindole (DAPI) from Cell Signaling Technology (Cat #4083S) for 15 min in the dark at 30 °C in the microaerobic chamber. FM 1-43 (Life Technologies Corporation Cat #T3163) was applied in the same way when required. Cells were then centrifuged at 10,000 x g for 3 min and washed 3 times with 100 µL fresh MG medium for 10 min in the dark at 30 °C in the microaerobic chamber. After washing, cells were resuspended in 10 µL fresh MG medium and 0.8 µL cell mixture was added to a slide and sealed under a coverslip using nail polish to reduce drying. Slides were imaged at 1000x magnification using the QImaging Retiga 1350ex camera in a Zeiss Axioimager M2 fluorescence microscope. Localization of proteins was quantified using the ImageJ Cell Counter plugin to categorize the localization in each cell into one of several categories including diffuse, foci, membrane, and chain aligned. Image file names were obscured using the ImageJ Randomizer macro for unbiased counting.

### 3D Structured illumination fluorescence microscopy and image analysis

Cells were prepared for fluorescence microscopy above and imaged using the Plan-APOCHROMAT 100x/1.46 objective lens of a Carl Zeiss Elyra PS.1 structured illumination microscope. Lasers at 405, 488, 561, and 642 nm wavelengths were used to excite DAPI, GFP, HaloTag ligand JF549, and HaloTag ligand JF646, respectively. Images were acquired using Zeiss ZEN software and processed using Imaris software (Bitplane).

### Pulse-chase analysis

To study Mms6 localization under changing iron conditions, we applied pulse-chase analysis using magnetosome proteins fused with HaloTag. HaloTag binds covalently and irreversibly to fluorescent ligands, allowing the tracking of a specific protein pool. For pulse-chase analysis, stock cultures were passaged into 10 mL fresh MG medium and grown in iron starvation conditions for 2 d in tubes washed with oxalic acid to remove residual iron. This process was repeated twice to ensure cells could not biomineralize. Then, 3 tubes of 10 mL AMB-1 cells per strain were grown in MG medium to early exponential phase (OD400 0.05-0.08) under iron starvation conditions. Cmag was assessed as described above for each culture. Cultures were pelleted by centrifugation at 10,000 x g for 3 min in an anaerobic chamber and resuspended with 500 nM HaloTag 549 pulse ligand and incubated in anaerobic MG medium for 60 min in the dark at 30 °C. Cells were centrifuged at 10,000 x g for 3 min. Cells were washed 3 times with 100 µL fresh, anaerobic MG medium for 10 min in the dark at 30 °C. Anaerobic MG medium was used to resuspend the cell pellets and the cell mixtures were inoculated into sealed anaerobic Balch tubes and incubated in the dark at 30 °C. 20µM Ferric malate was added to induce biomineralization, and OD400 and Cmag was tracked. 1 h before time point collection, cultures were centrifuged at 10,000 x g for 3 min. Cell pellets were resuspended with 500 nM HaloTag 646 (Promega Cat #GA1120) chase ligand in anaerobic MG medium for 45 min in the dark at 30 °C. 1.4 µM 4’,6-diamidino-2-phenylindole (DAPI) was added to cells, cells were mixed, and incubation continued for an additional 15 min. Cells were then centrifuged at 10,000 x g for 3 min and washed 3 times with 500 µL fresh MG medium. After washing, cells were resuspended in 10 µL fresh MG medium and 0.8 µL cell mixture was added to a slide and sealed under a coverslip using nail polish to reduce drying. Slides were imaged at 1000x magnification using the QImaging Retiga 1350ex camera in a Zeiss Axioimager M2 fluorescence microscope. Localization of proteins was quantified using the ImageJ Cell Counter plugin to categorize the localization in each cell into one of several categories including diffuse, foci, and chain aligned. Image file names were obscured using the ImageJ Randomizer macro for unbiased counting.

### Cellular fractionation

AMB-1 cells were grown in 50 mL MG medium at 30 °C in a microaerobic chamber maintaining 10% atmospheric oxygen. Cells were then diluted 1:100 into 1.5 L MG medium and grown for 2 d. The 1.5 L cultures were centrifuged at 8,000 xg for 20 min at 4 °C.

Pellets were resuspended in 1 mL Buffer A (10 mM Tris-HCl, pH 8.0, 50mM NaCl, 1mM EDTA). Pepstatin and Leupeptin were each added to a final concentration of 2 µg/ml and 2 mM PMSF was added. To lyse cells, 0.5 mg/mL lysozyme was added and samples were incubated at room temperature for 15 min. After lysis, 3mL Buffer B (20mM HEPES-KOH pH 7.5, 50mM NaCl, 1.25mM CaCl2) was added along with 2 mM DTT and 5 µg/mL DNAse I and lysates were rocked at 4 °C for 15 min. To separate soluble and insoluble cell fractions, samples were ultracentrifuged at 160,000 xg for 2 h at 4 °C in ultracentrifuge tubes (Beckman Coulter Cat #328874). The resulting pellet contained the insoluble AMB-1 cell fraction and the supernatant contained the soluble fraction. In fractionations done with Igepal CA-630 (Spectrum Chemicals Cat #I1112-100 ML), also known as Nonidet P-40 substitute or NP-40, 0.4% Igepal was added before ultracentrifugation and samples were kept on ice for 2 h and gently agitated every 30 min to mix.

Cell fractions were analyzed by SDS-PAGE. Briefly, 2x Laemmli Sample Buffer (Bio-Rad) was mixed with each fraction. After heating fractions for 15 min at 95 °C, proteins were resolved by electrophoresis through 12% agarose polyacrylamide gels and then transferred to PVDF membranes (Bio-Rad Cat #1620175) by electroblotting. Protein detection was done using primary antibody anti-HaloTag monoclonal antibody (1:1,000 dilution, Promega), primary antibody anti-MamE polyclonal antibody (1:3,000 dilution, produced by ProSci Inc), secondary antibody F(ab’)2-goat anti-mouse IgG (H+L) HRP-conjugate (1:5,000 dilution, Invitrogen), and secondary antibody goat anti-rabbit IgG (H+L)-HRP-conjugate (1:10,000 dilution, Bio-Rad). Image lab (Bio-Rad) software was used to take images of blots.

### Statistics and reproducibility

The chi-square test of independence or fisher’s exact test were used to assess significant differences in localization pattern distribution between samples. Mann-Whitney U test is a non-parametric test used to compare outcomes between two independent groups. The statistical tests were performed in RStudio using R version 4.2.2.

### Protein structure prediction

SignalP 5.0 was used to detect signal peptides in Mms6 and other magnetosome proteins. TMHMM 2.0 and TMPred were used to detect transmembrane regions. Phyre2 and CCTOP were used for membrane topology predictions.

---

**Supplementary Figure 1.**
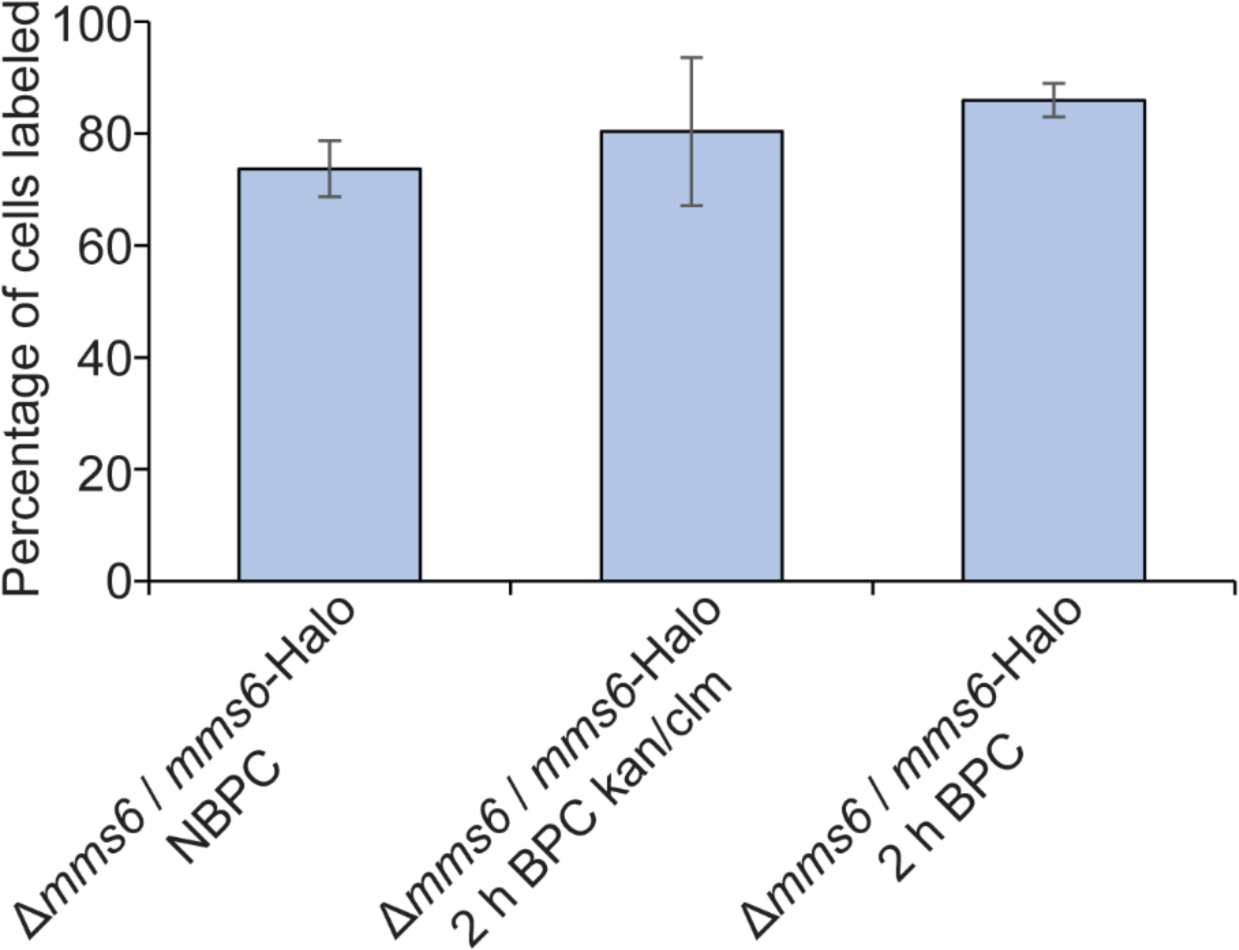
Percentage of cells labeled with Mms6-Halo fluorescence before and during relocalization time course with and without antibiotics. NBPC *n* = 2881 DAPI labeled cells, 2 h BPC kan/clm *n* = 1315 DAPI labeled cells, 2 h BPC *n* = 2822 DAPI labeled cells.

**Supplementary Figure 2.**
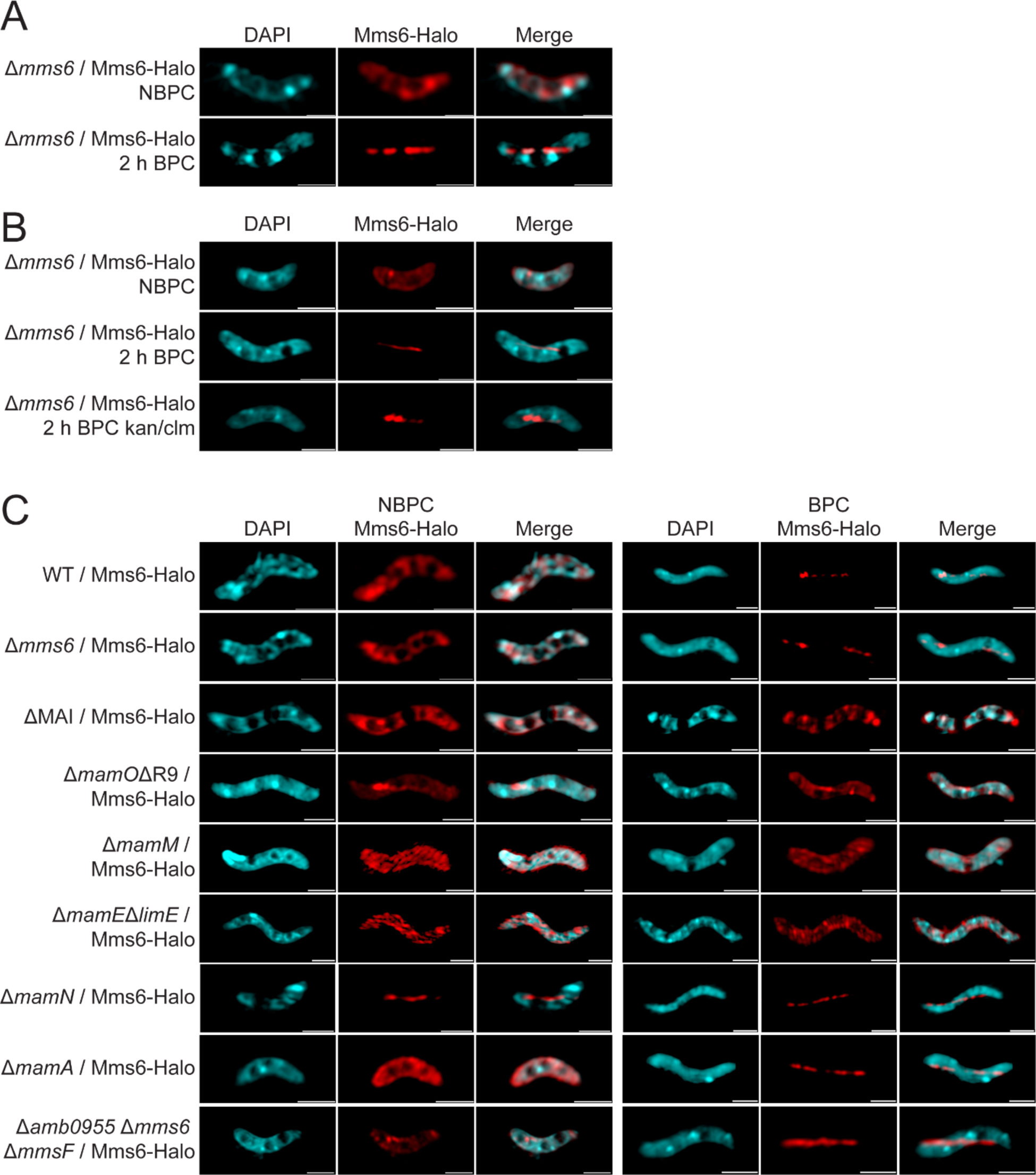
Representative super resolution 3D Structured Illumination Microscopy (SIM) images of cells expressing Mms6-Halo. (A) Representative images of WT cells expressing Mms6-Halo from biomineralization time course. WT cells grown under standard growth conditions expressing either GFP-MmsF or GFP-Mms6_NTD_MmsF shown in green and DAPI shown in blue. Sample labeled “kan/clm” was given 700 µg/ml kanamycin and 400 µg/ml chloramphenicol to prevent the synthesis of new Mms6-Halo during and after the one-hour labeling step. Scale bars = 1 µm. (B) Representative images of cells of different mutant backgrounds expressing Mms6-Halo in red and DAPI shown in blue. Scale bars = 1 µm.

**Supplementary Figure 3.**
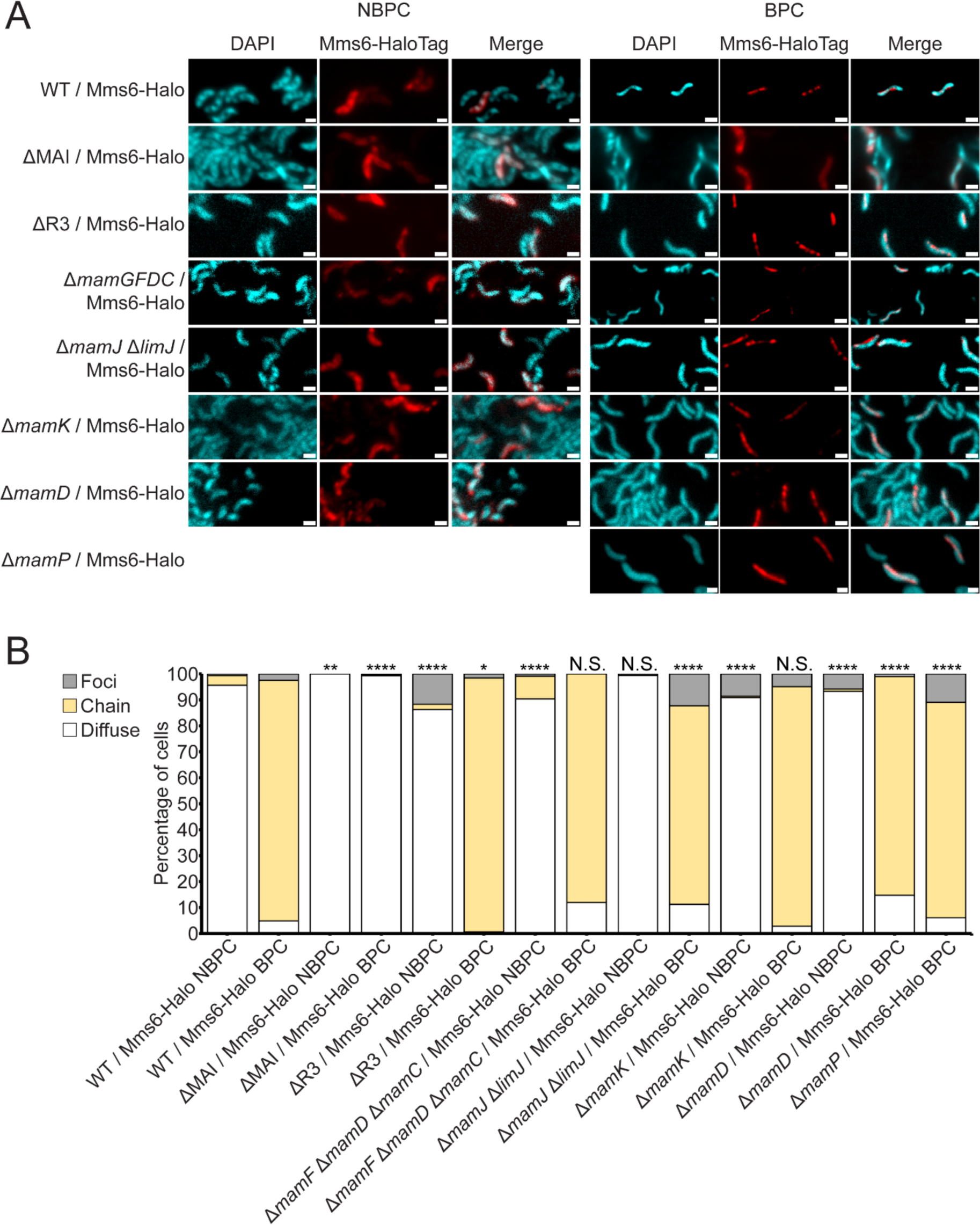
Mms6-Halo localization in further MAI protein deletion backgrounds (A) Representative fluorescence microscopy images of AMB-1 with different genetic backgrounds expressing Mms6-Halo and grown in standard conditions. JF549 HaloTag ligand fluorescence is shown in red and DAPI in blue. Scale bars = 1 µm. (B) Blind quantification of localization patterns of Mms6-Halo. *P* values were calculated by Fisher’s exact test comparing given dataset to WT / *mms6*-Halo (N.S. no significant difference *P* > .01) (* = *P* < .01) (** = *P* < 10^-3^) (**** = *P* < 10^-5^). WT / *mms6*-Halo NBPC *n* = 1010 cells, WT / *mms6*-Halo BPC *n* = 477 cells, ΔMAI / *mms6*-Halo NBPC *n* = 262 cells, ΔMAI / *mms6*-Halo BPC *n* = 400 cells, ΔR3 / *mms6*-Halo NBPC *n* = 197 cells, ΔR3 / *mms6*-Halo BPC *n* = 316 cells, Δ*mamF* Δ*mamD* Δ*mamC* / *mms6*-Halo NBPC *n* = 996 cells, Δ*mamF* Δ*mamD* Δ*mamC* / *mms6*-Halo BPC *n* = 92 cells, Δ*mamJ* Δ*limJ* / *mms6*-Halo NBPC *n* = 160 cells, Δ*mamJ* Δ*limJ* / *mms6*-Halo BPC *n* = 277 cells, Δ*mamK* / *mms6*-Halo NBPC *n* = 175 cells, Δ*mamK* / *mms6*-Halo BPC *n* = 142 cells, Δ*mamD* / *mms6*-Halo NBPC *n* = 223 cells, Δ*mamD* / *mms6*-Halo BPC *n* = 197 cells.

**Supplementary Figure 4.**
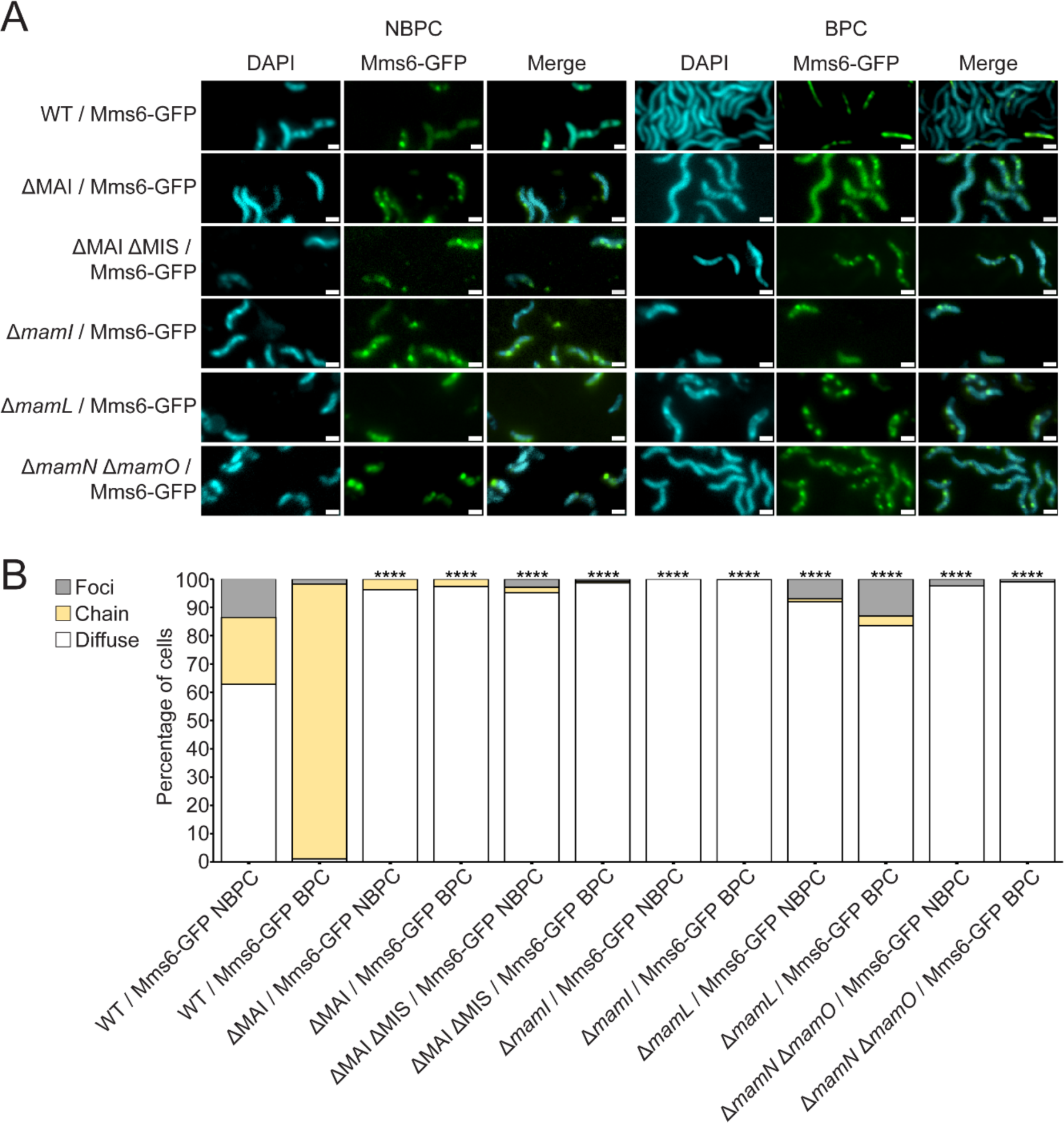
Mms6-GFP localization in further MAI protein deletion backgrounds (A) Representative fluorescence microscopy images of AMB-1 with different genetic backgrounds expressing Mms6-GFP and grown in standard conditions. GFP is shown in green and DAPI in blue. Scale bars = 1 µm. (B) Blind quantification of localization patterns of Mms6-GFP. *P* values were calculated by Fisher’s exact test comparing given dataset to WT / *mms6*-GFP (**** = *P* < 10^-5^). WT / *mms6*-GFP NBPC *n* = 199 cells, WT / *mms6*-GFP BPC *n* = 282 cells, ΔMAI / *mms6*-GFP NBPC *n* = 356 cells, ΔMAI / *mms6*-GFP BPC *n* = 1070 cells, ΔMAI ΔMIS / *mms6*-GFP NBPC *n* = 104 cells, ΔMAI ΔMIS / *mms6*-GFP BPC *n* = 540 cells, Δ*mamI* / *mms6*-GFP NBPC *n* = 72 cells, Δ*mamI* / *mms6*-GFP BPC *n* = 77 cells, Δ*mamL* / *mms6*-GFP NBPC *n* = 487 cells, Δ*mamL* / *mms6*-GFP BPC *n* = 207 cells, Δ*mamN* Δ*mamO* / *mms6*-GFP NBPC *n* = 82 cells, Δ*mamN* Δ*mamO* / *mms6*-GFP BPC *n* = 715 cells.

**Supplementary Figure 5.**
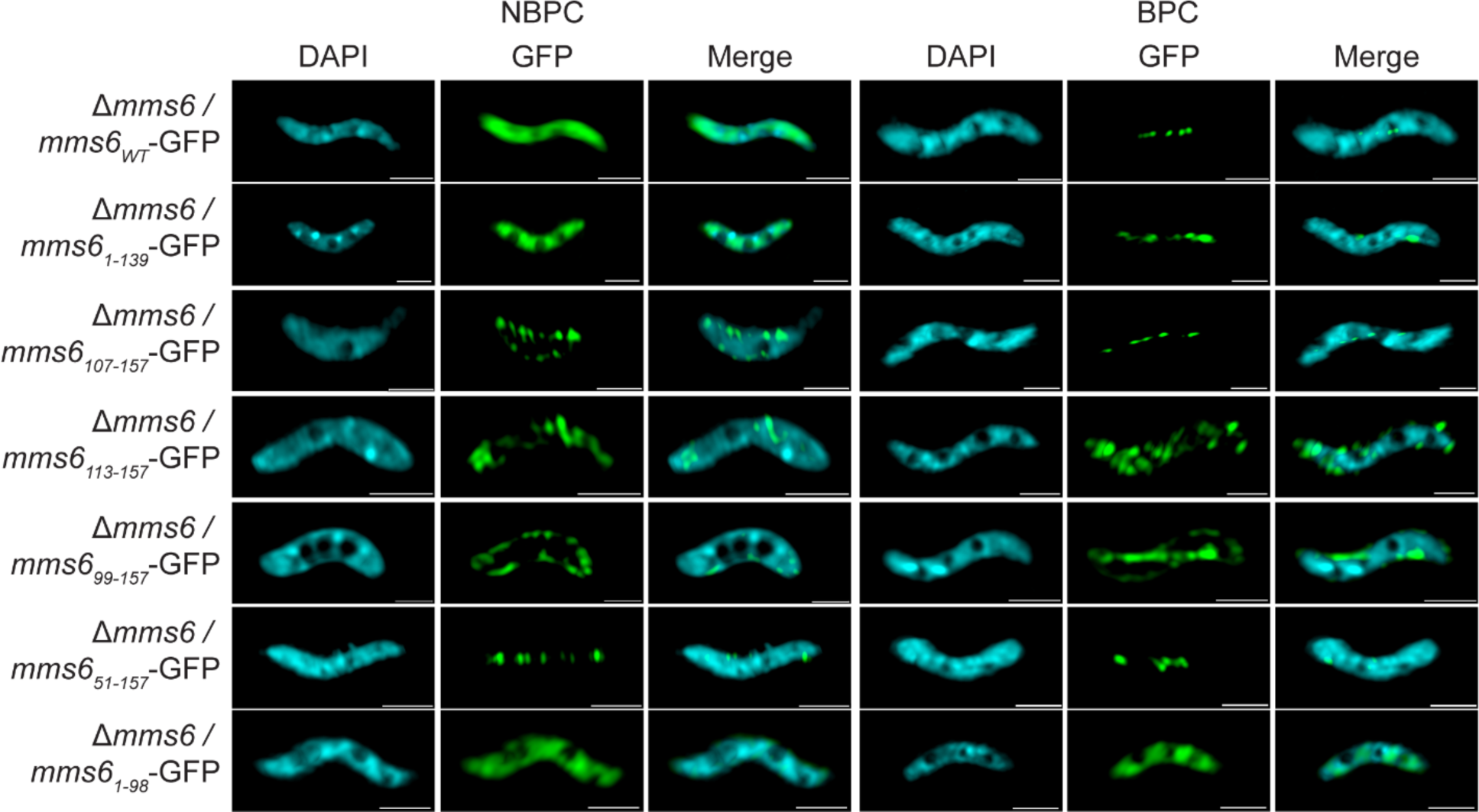
Representative super resolution 3D Structured Illumination Microscopy (SIM) images of cells expressing Mms6-GFP. Representative images of Δ*mms6* mutant cells expressing Mms6-GFP or a mutant Mms6 protein tagged C-terminally with GFP. GFP is shown in green and DAPI in blue. Scale bars = 1 µm.

**Supplementary Figure 6.**
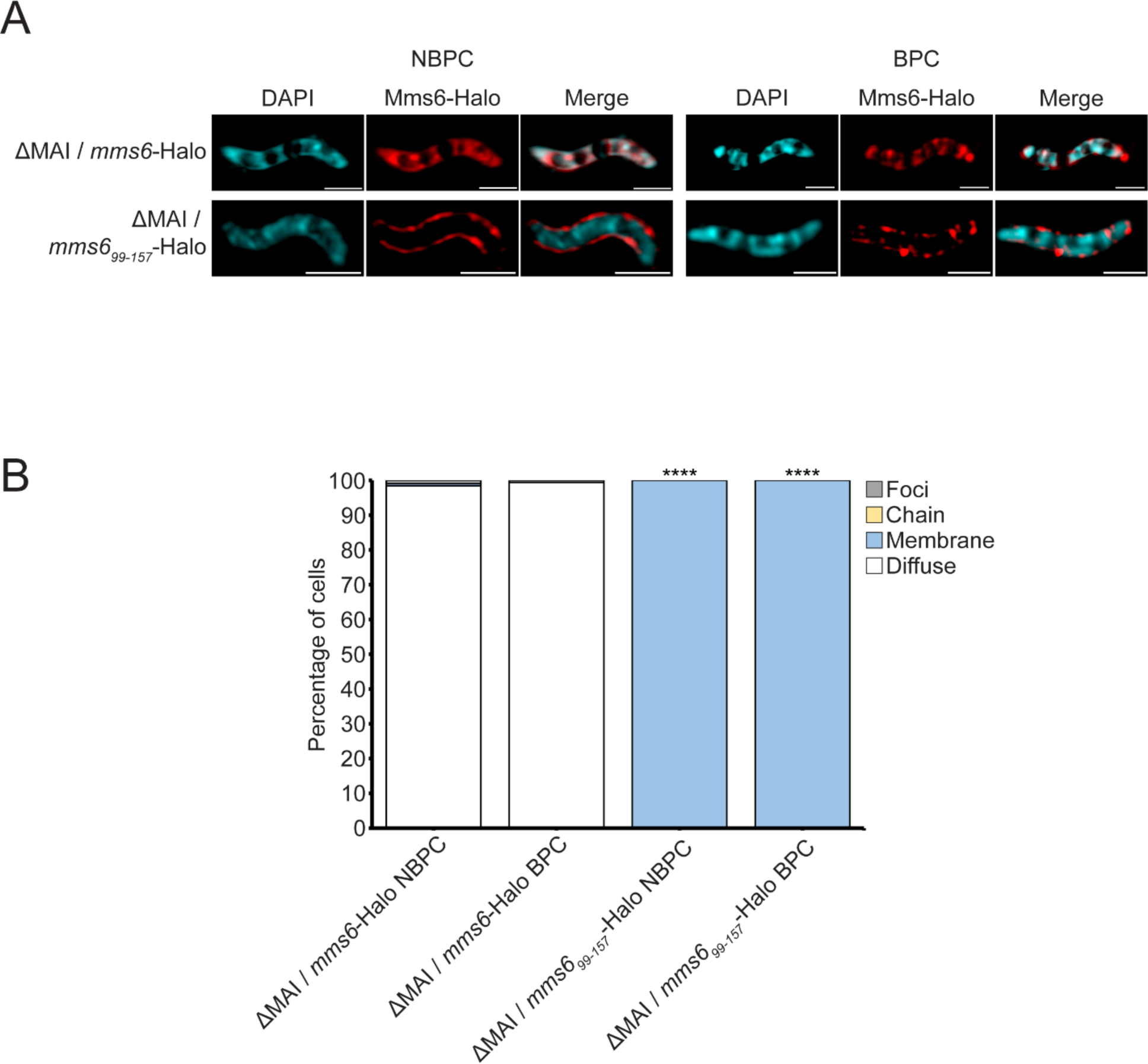
Mms6_99-157_-Halo does not require other magnetosome proteins to translocate into membranes. (A) Representative images of ΔMAI mutant cells expressing Mms6-Halo or Mms6_99-157_-Halo shown in red. DAPI counterstain is shown in blue. Scale bars = 1 µm. (B) Blind quantification of localization patterns of given protein. *P* values were calculated by Fisher’s exact test comparing given dataset to ΔMAI / *mms6*-Halo grown in matching biomineralization condition (**** = *P* < 10^-5^). ΔMAI / *mms6*-Halo NBPC n = 318 cells, ΔMAI / *mms6*-Halo BPC n = 174 cells, ΔMAI / *mms6*_99-157_-Halo NBPC n = 82 cells, ΔMAI / *mms6*_99-157_-Halo BPC n = 23 cells.

**Supplementary Figure 7.**
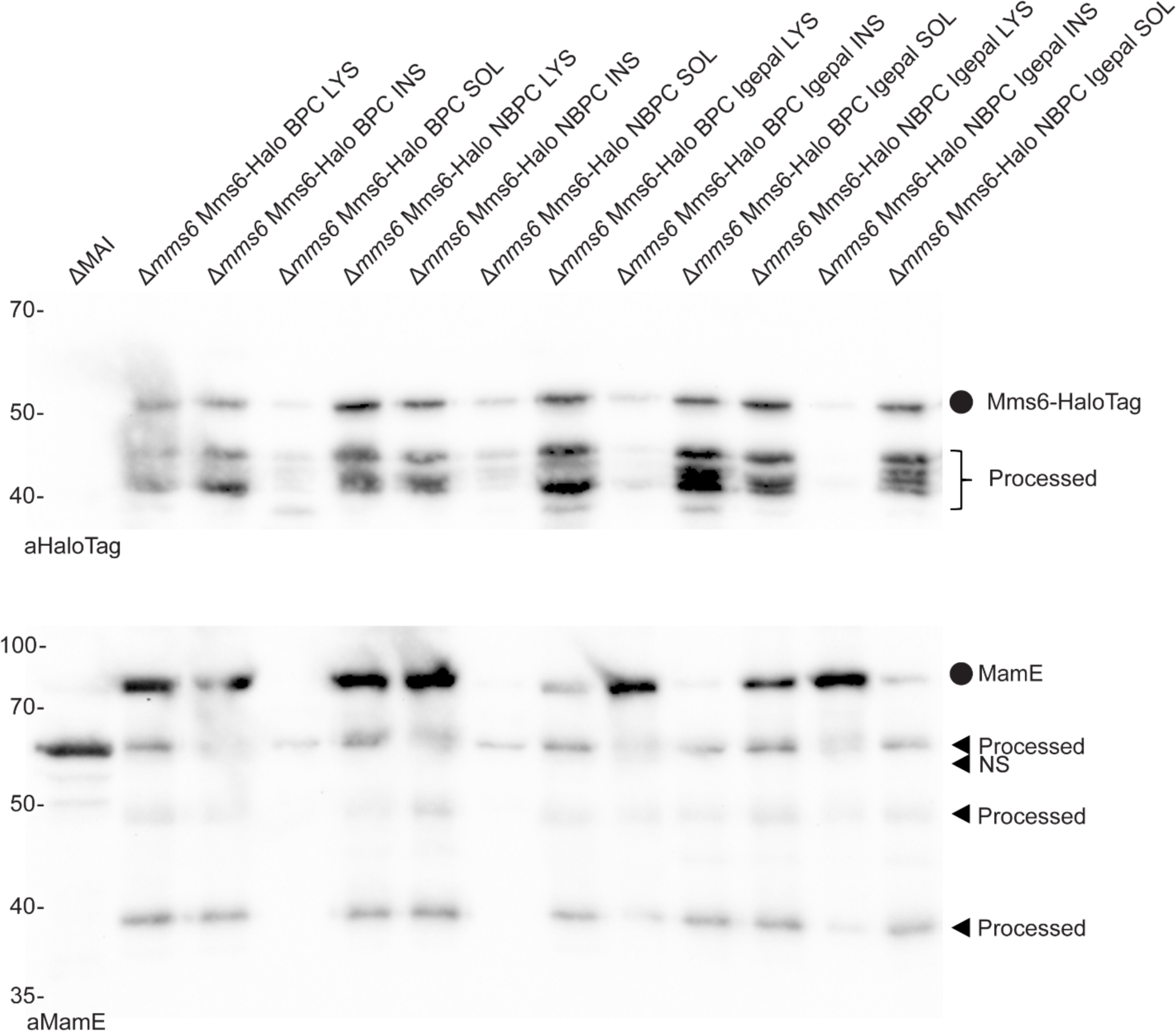
Mms6 is weakly associated with insoluble cell contents. Immunoblotting analysis of cell fractions after fractionation of cells expressing Mms6-Halo. HaloTag and MamE were probed for in the whole-cell lysate (LYS) before centrifugation, as well as in insoluble (INS) and soluble (SOL) fractions afterwards. The cell fractionation was performed with and without 0.4% Igepal. Full length protein bands are marked with circles, and both processed fragments and non-specific bands (NS) are marked with arrows.

**Supplementary Figure 8.**
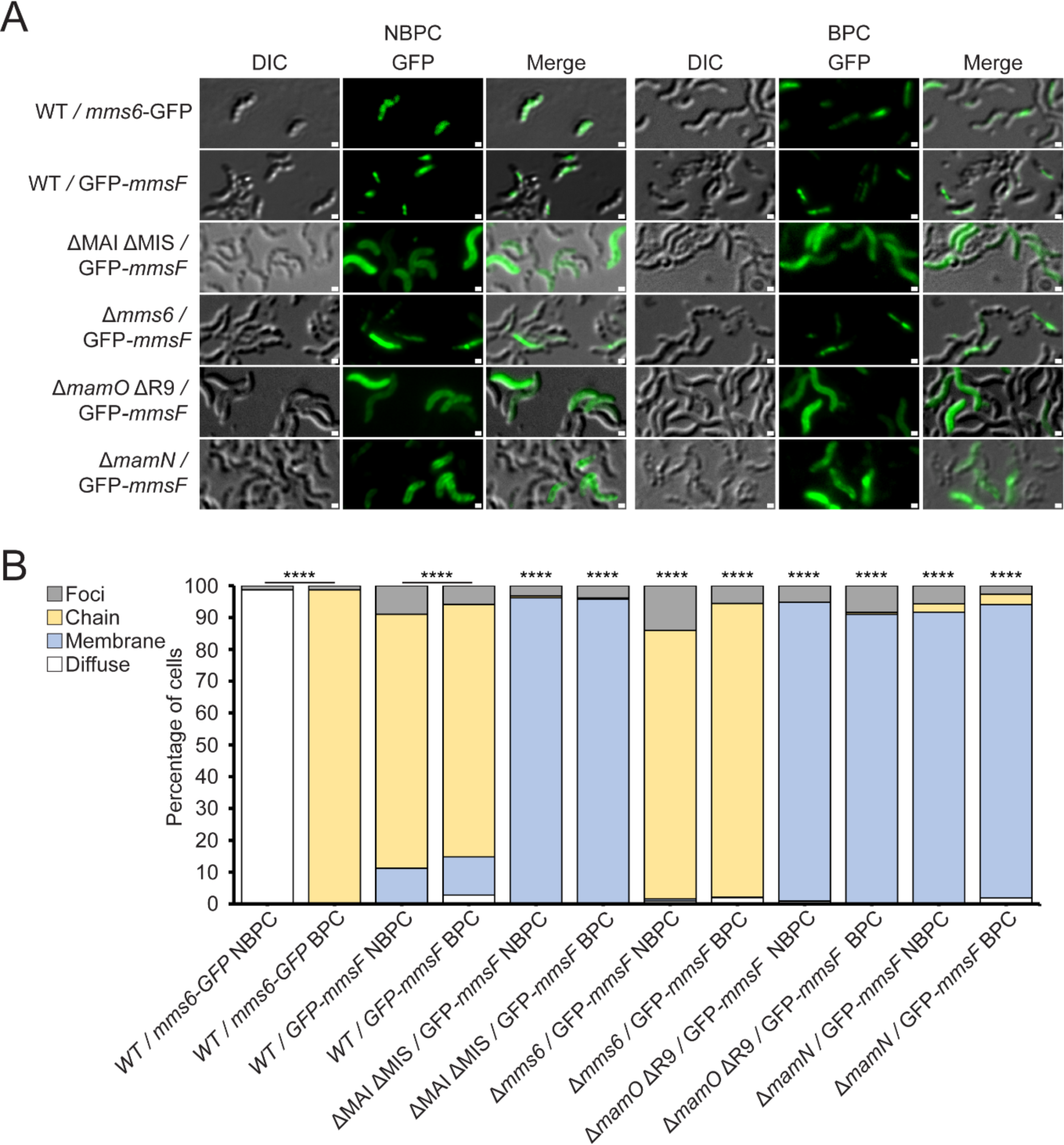
MmsF requires *mamN* and *mamO* for magnetosome localization. (A) Representative fluorescence microscopy images of WT or mutant AMB-1 cells grown under standard growth conditions expressing Mms6 or MmsF GFP fusions. GFP is shown in green and transmitted light (TL) is displayed to show outlines of AMB-1 cells. Scale bars = 1 µm. (B) Blind quantification of localization patterns of GFP tagged magnetosome proteins *in vivo* expressed in either WT or *Δmms6* cells. The Y-axis represents percentage of total cell count with indicated protein fluorescence pattern. *P* values were calculated by Fisher’s exact test comparing indicated datasets (**** *P* < 10^-5^). WT / *mms6*-GFP NBPC *n* = 1074 cells, WT / *mms6*-GFP BPC *n* = 1317 cells, WT / GFP-*mmsF* NBPC *n* = 2412 cells, WT / GFP-*mmsF* BPC *n* = 1295 cells, ΔMAI ΔMIS / GFP-*mmsF* NBPC *n* = 1034 cells, ΔMAI ΔMIS / GFP-*mmsF* BPC *n* = 519 cells, *Δmms6* / GFP-*mmsF* NBPC *n* = 497 cells, *Δmms6* / GFP-*mmsF* BPC *n* = 1062 cells, Δ*mamO* ΔR9 / GFP-*mmsF* NBPC *n* = 478 cells, Δ*mamO* ΔR9 / GFP-*mmsF* BPC *n* = 91 cells, Δ*mamN* / GFP-*mmsF* NBPC *n* = 421 cells, Δ*mamN* / GFP-*mmsF* BPC *n* = 524 cells.

**Supplementary Figure 9.**
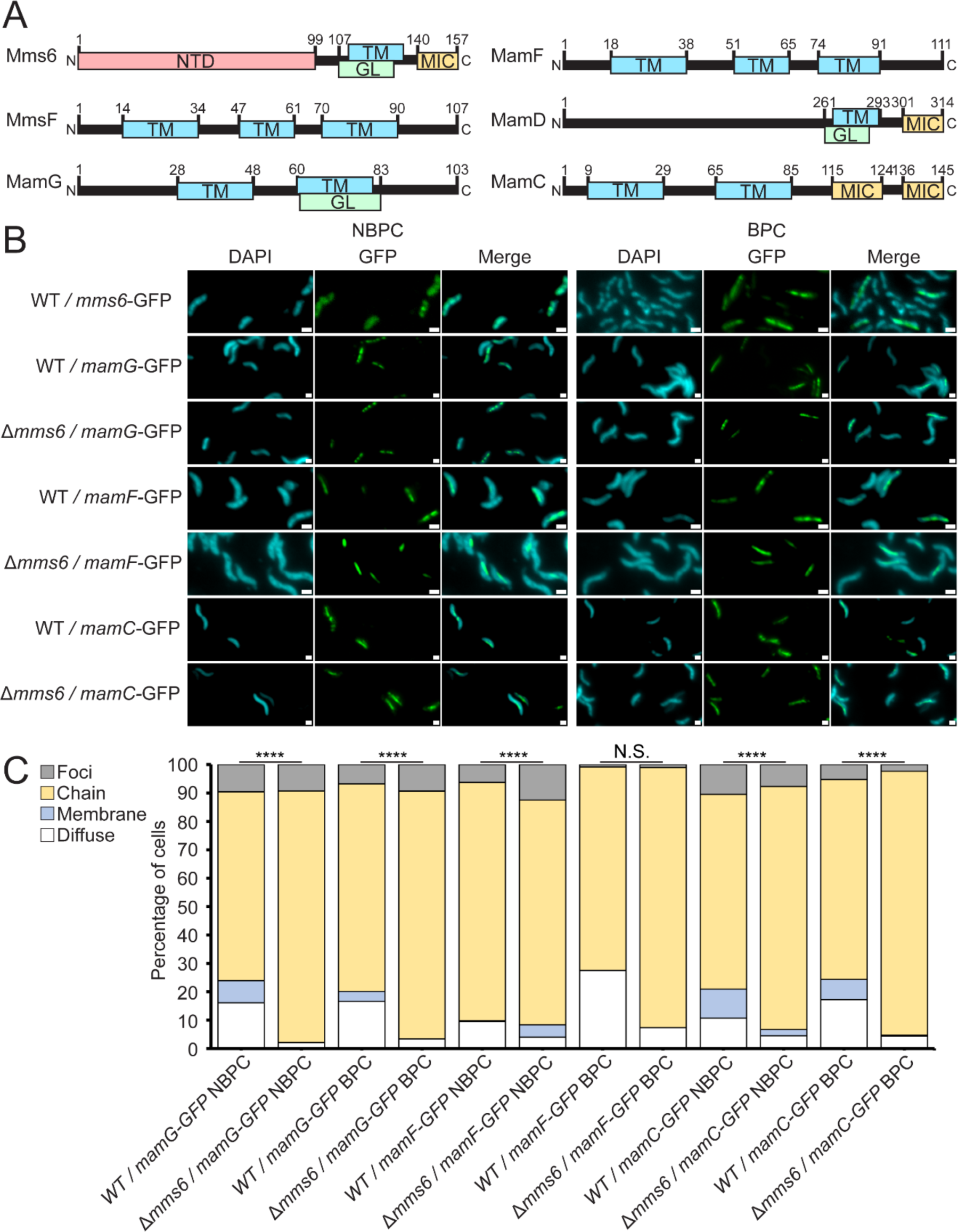
Magnetite shaping proteins do not require *mms6* for magnetosome localization. (A) Structural features of several magnetosome proteins implicated in crystal maturation. N-terminal domains (NTD), glycine leucine repeat domains (GL), transmembrane domains (TM), magnetite-interacting component (MIC). (B) Representative fluorescence microscopy images of WT or Δ*mms6* AMB-1 cells grown under standard growth conditions expressing magnetosome protein GFP fusions. GFP is shown in green, and DAPI in blue. Scale bars = 1 µm. (C) Blind quantification of localization patterns of GFP tagged magnetosome proteins *in vivo* expressed in WT or Δ*mms6* cells. The Y-axis represents percentage of total cell count with indicated protein fluorescence pattern. *P* values were calculated by Fisher’s exact test comparing indicated datasets (N.S. no significant difference *P* > .01) (**** *P* < 10^-5^). WT / *mamG*-GFP NBPC *n* = 1116 cells, *Δmms6* / *mamG*-GFP NBPC *n* = 1440 cells, WT / *mamG*-GFP BPC *n* = 367 cells, *Δmms6* / *mamG*-GFP BPC *n* = 1560 cells, WT / *mamF*-GFP NBPC *n* = 1238 cells, *Δmms6* / *mamF*-GFP NBPC *n* = 1583 cells, WT / *mamF*-GFP BPC *n* = 1784 cells, *Δmms6* / *mamF*-GFP BPC *n* = 2381 cells, WT / *mamC*-GFP NBPC *n* = 372 cells, *Δmms6* / *mamC*-GFP NBPC *n* = 793 cells, WT / *mamC*-GFP BPC *n* = 1253 cells, *Δmms6* / *mamC*-GFP BPC *n* = 1881 cells.

**Supplementary Table 1.**
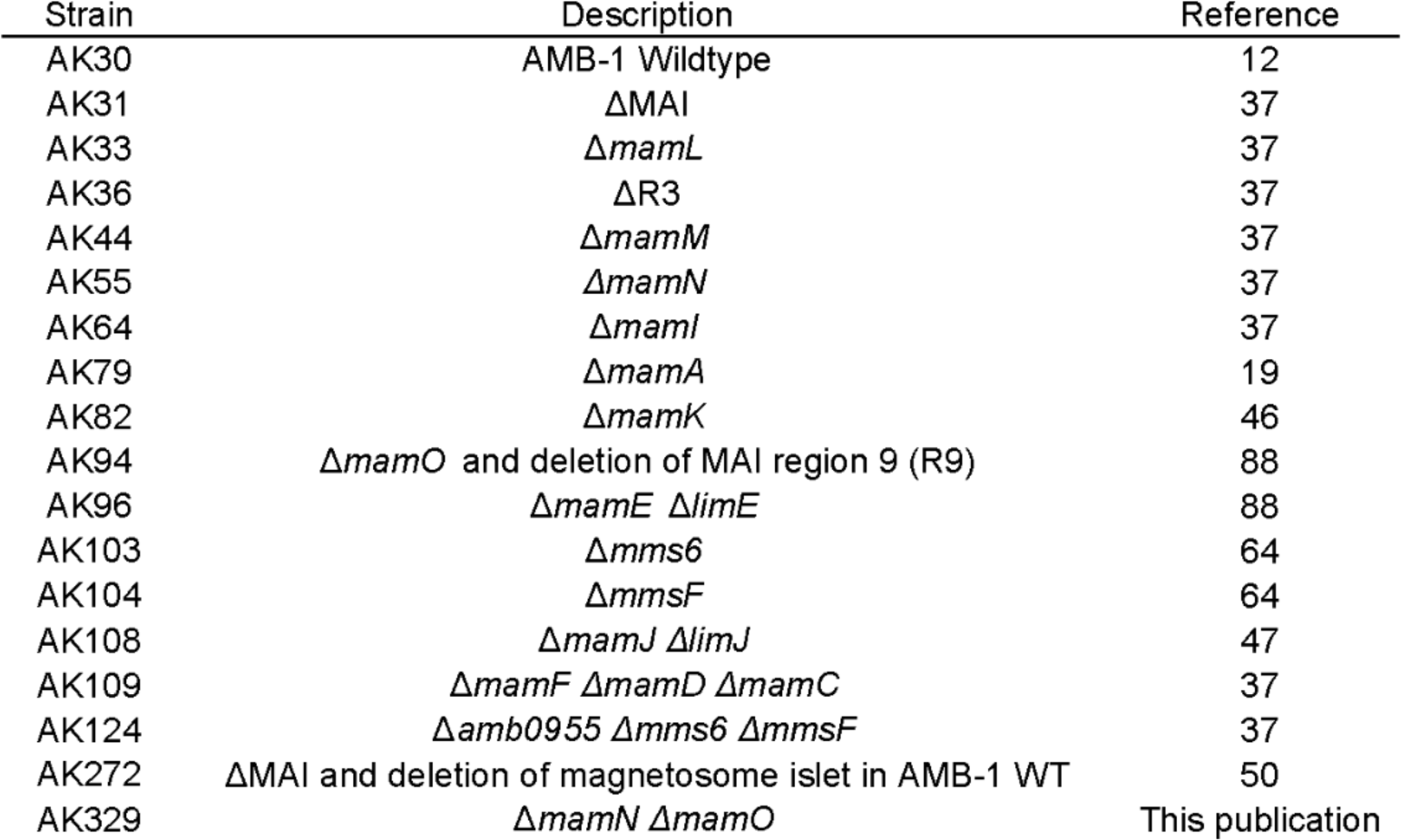
Strains used in this study.

| Strain | Description | Reference |
| --- | --- | --- |
| AK30 | AMB-1 Wildtype | 12 |
| AK31 | ΔMAI | 37 |
| AK33 | ΔmamL | 37 |
| AK36 | ΔR3 | 37 |
| AK44 | ΔmamM | 37 |
| AK55 | ΔmamN | 37 |
| AK64 | ΔmamI | 37 |
| AK79 | ΔmamA | 19 |
| AK82 | ΔmamK | 46 |
| AK94 | ΔmamO and deletion of MAI region 9 (R9) | 88 |
| AK96 | ΔmamE ΔlimE | 88 |
| AK103 | Δmms6 | 64 |
| AK104 | ΔmmsF | 64 |
| AK108 | ΔmamJ ΔlimJ | 47 |
| AK109 | ΔmamF ΔmamD ΔmamC | 37 |
| AK124 | Δamb0955 Δmms6 ΔmmsF | 37 |
| AK272 | ΔMAI and deletion of magnetosome islet in AMB-1 WT | 50 |
| AK329 | ΔmamN ΔmamO | This publication |

**Supplementary Table 2.** Plasmids used in this study.

| Plasmid | Description | Plasmid backbone |
| --- | --- | --- |
| pAK272 | Ptac-*mamI* -GFP | pAK22 |
| pAK452 | Ptac-*mamC* -GFP | pAK22 |
| pAK454 | Ptac-*mamF* -GFP | pAK22 |
| pAK532 | Ptac-GFP-*mmsF* | pAK22 |
| pAK976 | Ptac-mHaloTag | pAK22 |
| pAK1101 | Ptac-*mms6* -HaloTag | pAK22 |
| pAK1102 | Ptac-*mms6* -GFP | pAK22 |
| pAK1204 | Ptac-*mamD* -GFP | pAK22 |
| pAK1440 | Ptac-*mamG* -GFP | pAK22 |
| pAK1441 | Ptac-*mms6*<sub>51-157</sub> -GFP | pAK1102 |
| pAK1443 | Ptac-*mms6*<sub>1-139</sub> -GFP | pAK1102 |
| pAK1444 | Ptac-*mms6*<sub>1-98</sub> -GFP | pAK1102 |
| pAK1445 | Ptac-*mms6*<sub>107-157</sub> -GFP | pAK1102 |
| pAK1446 | Ptac-*mms6*<sub>113-157</sub> -GFP | pAK1102 |
| pAK1447 | Ptac-GFP-*mms6*<sub>NTD</sub>*mmsF* | pAK532 |
| pAK1456 | Ptac-*mms6*<sub>99-157</sub> -GFP | pAK1102 |

**Supplementary Table 3.**
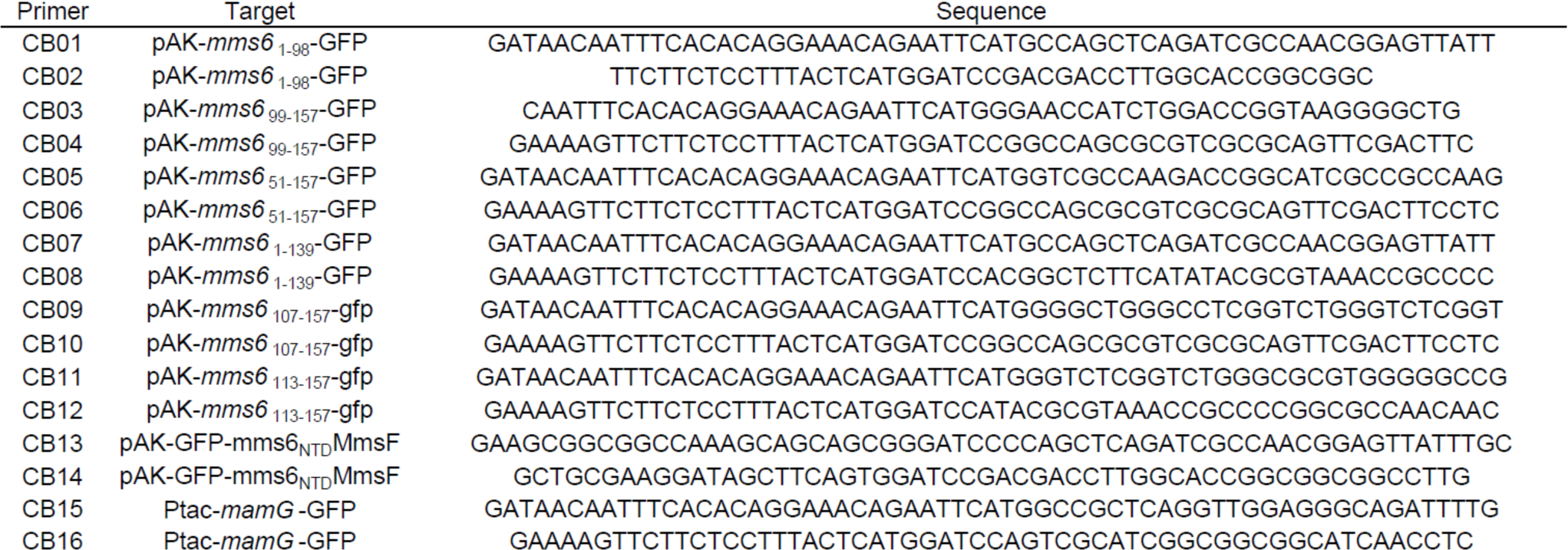
Primers used in this study.

**Supplementary Table 4.** Statistical tests using the Chi-square test of independence. The Chi-square test of independence tests the hypothesis that two variables are unrelated. Cramer’s *V* is an effect size measurement that measures how strongly two categorical fields are associated.

| Figure | Dataset/strain | Compared datasets/strains | Chi-square test of independence |  | Cramer's V effect size |  |  |
| --- | --- | --- | --- | --- | --- | --- | --- |
|  |  |  | P-value | Significance | V | df | Effect Size |
| Figure 1B | NBPC | 30 min BPC | 0.3517 | NS | 0.0422 | 3 | Negligible |
| | NBPC | 1 h BPC | $< 10^{-5}$ | **** | 0.3162 | 3 | Large |
| | NBPC | 1.5 h BPC | $< 10^{-5}$ | **** | 0.3496 | 3 | Large |
| | NBPC | 2 h BPC | $< 10^{-5}$ | **** | 0.4602 | 3 | Large |
| Figure 1I | NBPC | 2 h BPC | $< 10^{-5}$ | **** | 0.5654 | 2 | Large |
| Figure 2E | NBPC | 2 h BPC kan/clm | $< 10^{-5}$ | **** | 0.7124 | 3 | Large |
| | NBPC | 2 h BPC | $< 10^{-5}$ | **** | 0.6997 | 3 | Large |

**Supplementary Table 5.**
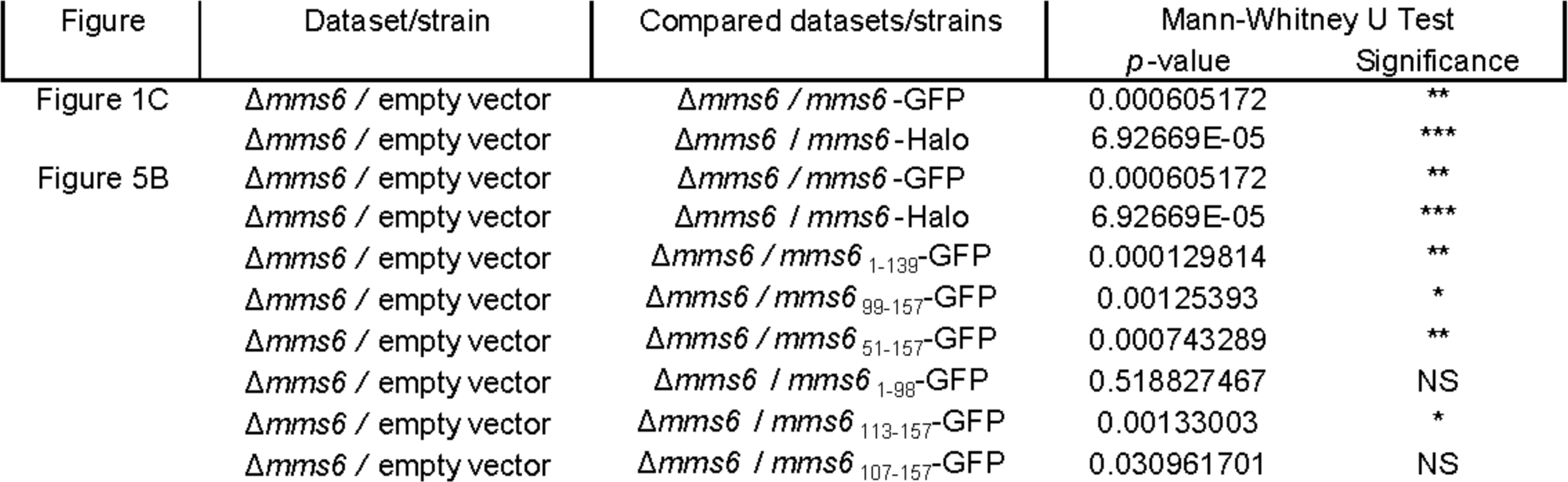
Statistical tests using the Mann Whitney *U* test. The Mann Whitney *U* test is a non-parametric test for the null hypothesis that the means of two populations are equal.

**Supplementary Table 6.**
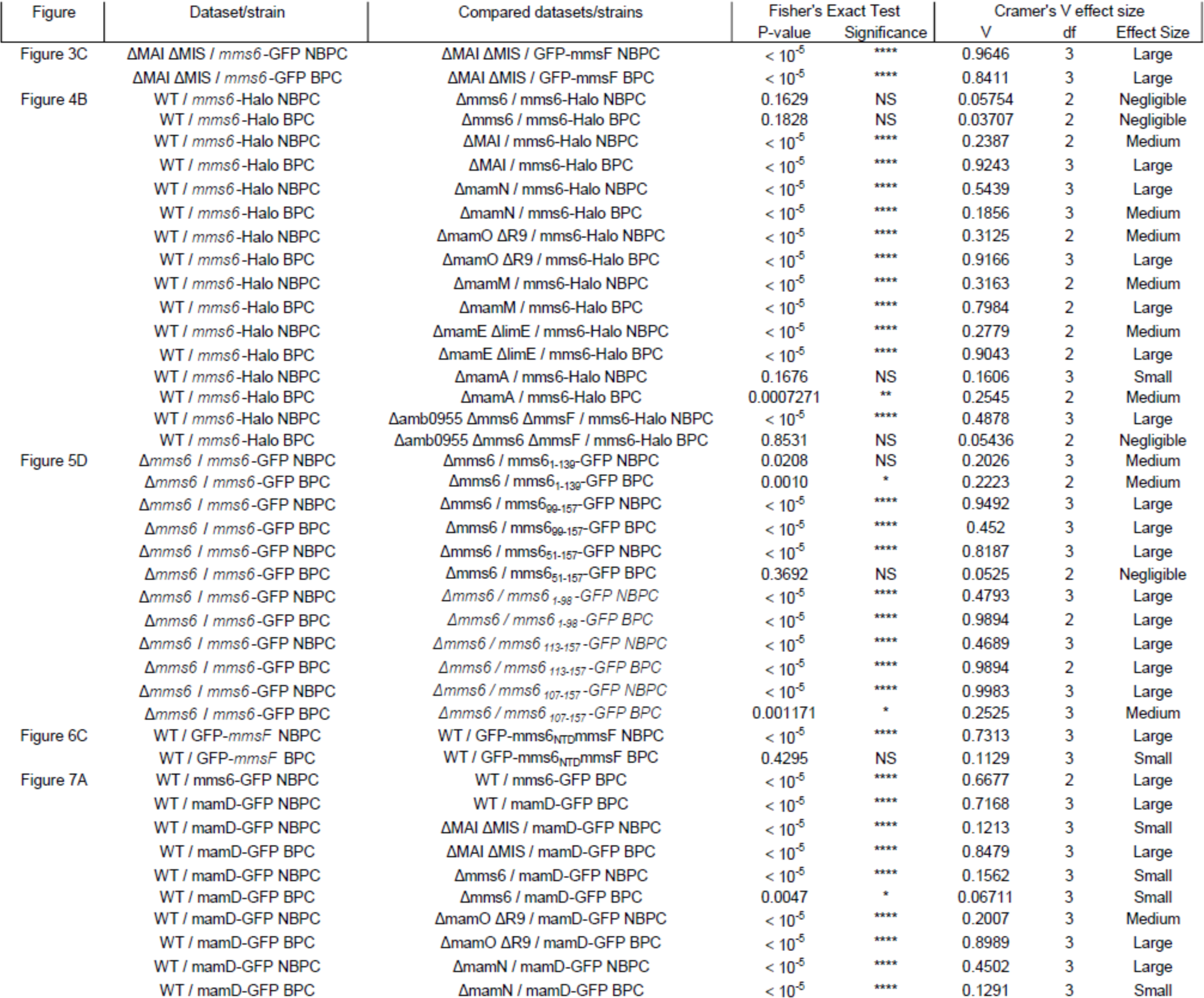
Statistical tests using the Fisher’s exact test. Fisher’s exact test tests the same hypothesis as the chi-squared test of independence but is more accurate for sample sizes under 500 and in cases where at least one sample has a value of zero. Cramer’s *V* is an effect size measurement that measures how strongly two categorical fields are associated.

**Supplementary Table 7.**
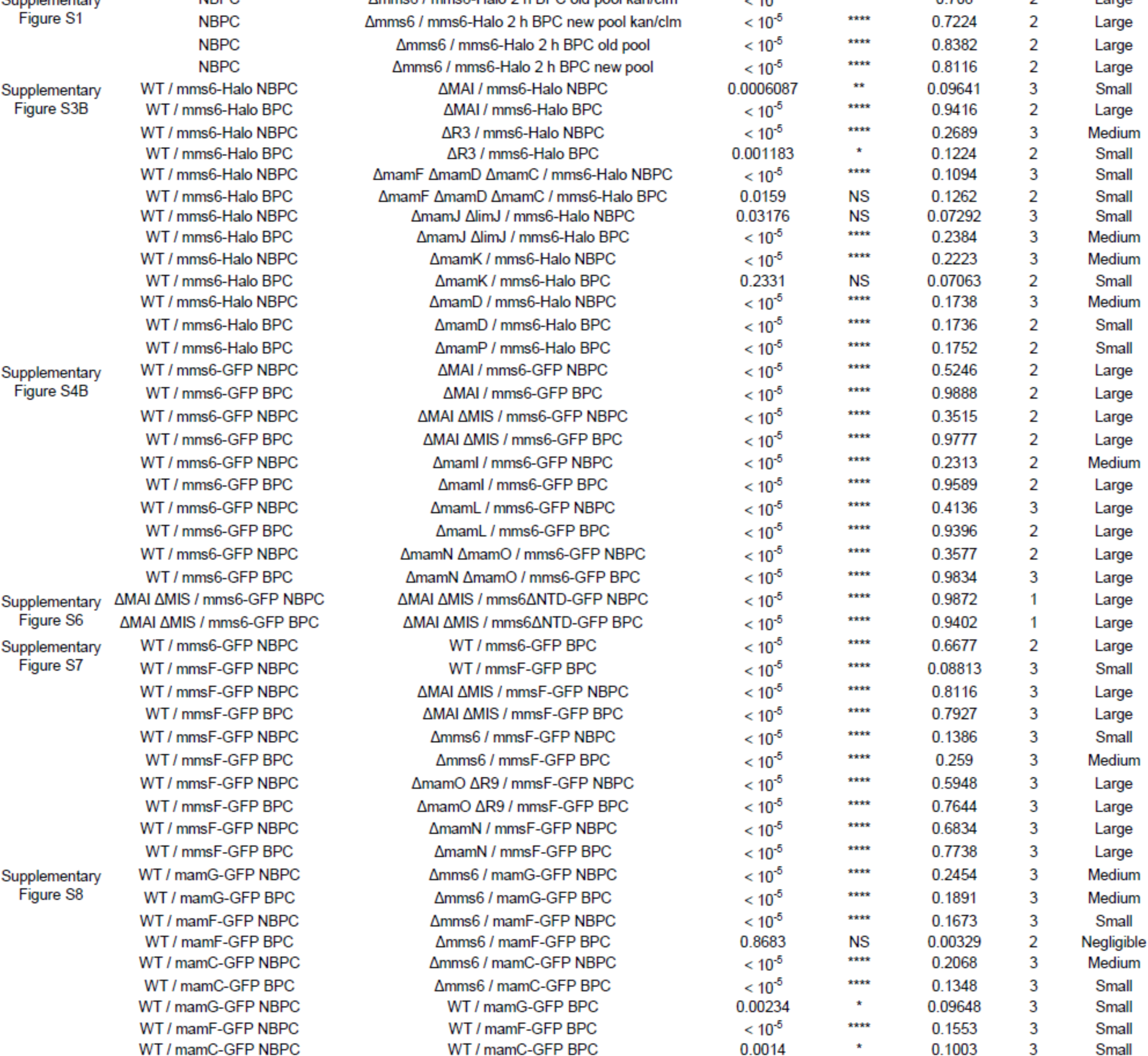
Statistical tests for supplementary figures. Fisher’s exact test tests the same hypothesis as the chi-squared test of independence but is more accurate for sample sizes under 500 and in cases where at least one sample has a value of zero. Cramer’s *V* is an effect size measurement that measures how strongly two categorical fields are associated.

